# FAM122A ensures cell cycle interphase progression and checkpoint control as a SLiM-dependent substrate-competitive inhibitor to the B55⍺/PP2A phosphatase

**DOI:** 10.1101/2023.03.06.531310

**Authors:** Jason S Wasserman, Bulat Faezov, Kishan R Patel, Alison N Kurimchak, Seren M Palacio, Holly Fowle, Brennan C McEwan, Qifang Xu, Ziran Zhao, Lauren Cressey, Neil Johnson, James S Duncan, Arminja N Kettenbach, Roland L Dunbrack, Xavier Graña

## Abstract

The Ser/Thr protein phosphatase 2A (PP2A) is a highly conserved collection of heterotrimeric holoenzymes responsible for the dephosphorylation of many regulated phosphoproteins. Substrate recognition and the integration of regulatory cues are mediated by B regulatory subunits that are complexed to the catalytic subunit (C) by a scaffold protein (A). PP2A/B55 substrate recruitment was thought to be mediated by charge-charge interactions between the surface of B55α and its substrates. Challenging this view, we recently discovered a conserved SLiM [**RK**]-**V**-x-x-[**VI**]-**R** in a range of proteins, including substrates such as the retinoblastoma-related protein p107 and TAU (Fowle *et al. eLife* 2021;10:e63181). Here we report the identification of this SLiM in FAM122A, an inhibitor of B55α/PP2A. This conserved SLiM is necessary for FAM122A binding to B55α *in vitro* and in cells. Computational structure prediction with *AlphaFold2* predicts an interaction consistent with the mutational and biochemical data and supports a mechanism whereby FAM122A uses the ‘SLiM’ in the form of a short α-helix to dock to the B55α top groove. In this model, FAM122A spatially constrains substrate access by occluding the catalytic subunit with a second α-helix immediately adjacent to helix 1. Consistently, FAM122A functions as a competitive inhibitor as it prevents binding of substrates in *in vitro* competition assays and the dephosphorylation of CDK substrates by B55α/PP2A in cell lysates. Ablation of FAM122A in human cell lines reduces the rate of proliferation, progression through cell cycle transitions and abrogates G1/S and intra-S phase cell cycle checkpoints. FAM122A-KO in HEK293 cells results in attenuation of CHK1 and CHK2 activation in response to replication stress. Overall, these data strongly suggest that FAM122A is a ‘SLiM’-dependent, substrate-competitive inhibitor of B55α/PP2A that suppresses multiple functions of B55α in the DNA damage response and in timely progression through the cell cycle interphase.

## Introduction

Protein phosphorylation is a reversible post-translational modification that plays critical roles in the regulation of many signaling pathways and cellular processes. It is estimated that two thirds of all human proteins are phosphorylated, with more than 98% of phosphorylation events occurring on serine and threonine residues^1, 2, 3^. Kinases catalyze the transfer of the γ-phosphate of ATP to phosphorylatable residues, while phosphatases hydrolyze the phosphate from phosphorylated residues. Members of the serine/threonine <u>p</u>hospho<u>p</u>rotein <u>p</u>hosphatase (PPP) family are responsible for the majority of dephosphorylation occurring in eukaryotic cells. Within this family, protein phosphatase 1 (PP1) and protein phosphatase 2A (PP2A) account for more than 90% of the total phosphatase activity^4, 5^. The PP2A Ser/Thr phosphatase forms multimeric complexes in cells. The heterodimeric “core enzyme” is made of a scaffold subunit (A) and a catalytic (C) subunit. For the formation of the heterotrimeric holoenzyme, the core dimer recruits a B subunit, which is the key determinant of substrate specificity for the holoenzyme reviewed in ^5, 6, 7, 8, 9^. There are four B families: B/B55, B’/B56, B’’/B72, and B’’’/B93; each consisting of three to five isoforms and splice variants. The mechanism of substrate recognition by serine/threonine phosphatases is typically dependent on short linear motifs (SLiMs) present in substrates, which enable a direct interaction with catalytic or regulatory subunits. For instance, the B56 family of B subunits bind substrates with a *LxxIxE* SLiM in Intrinsically <u>D</u>isordered <u>R</u>egions (IDRs)^10, 11, 12^. In contrast, PP2A/B55 substrate recruitment was thought to be mediated by charge-charge interactions between the surface of B55α and its substrates^13^. However, we have recently made an important discovery that challenges this view with significant implications for our understanding of PP2A biology, as PP2A/B55α is the most abundant PP2A holoenzyme. We identified a conserved SLiM [**RK**]-**V**-x-x-[**VI**]-**R** present in the intrinsically disordered regions (IDR) in a range of proteins, including substrates such as the retinoblastoma-related protein p107 and TAU^14^.

B55α, which is the most abundant isoform of the B/B55 family, has been implicated in the dephosphorylation of many substrates with functions in cellular signaling, the cell cycle and other cellular processes in post-mitotic cells^6^. PP2A/B55α holoenzymes are inactivated by a rapid switch at the G2/M transition that results in simultaneous activation of Cyclin B/CDK1. Inhibition of B55α is mediated by ARPP19 upon phosphorylation by a kinase called MASTL^15, 16^. We have shown that PP2A/B55α is also key for an interphase equilibrium that modulates the phosphorylation state of the retinoblastoma-related proteins^17, 18, 19^, and has also been implicated in checkpoint control^20, 21^. Our search for B55α binding proteins containing a degenerate B55α substrate SLiM identified FAM122A, a protein recently found to inhibit PP2A/B55α^22^. FAM122A has recently been shown to control activation of PP2A/B55α in response to massive replication stress caused by CHK1 inhibition in NSCLC cell lines, which activates the WEE1 kinase, a negative regulator of CDK1^23^. However, the mechanism by which FAM122A inhibits the PP2A/B55α holoenzyme is not well understood.

Using rigorous biochemical and molecular modeling approaches, we have generated a strong body of data supporting substrate competition as a mechanism for inhibition of PP2A/B55α. In addition, we report that ablation of FAM122A in cells inhibits proliferation, attenuates mitogenic signaling linked to the G0/G1 transition, and abrogates the G1/S checkpoint in response to nucleotide depletion by a mechanism associated with attenuation of CHK1 and CHK2 signaling leading to DNA damage and cell death. Therefore, FAM122A controls PP2A/B55α function using a decoy mechanism to block substrate access to the enzyme active site thereby controlling cell cycle transitions stimulated by mitogens and genotoxic stresses.

## Results

### A filtered proteome-wide search identifies FAM122A as a potential SLiM-containing PP2A/B55α interactor

Using extensive mutational analysis and competition assays, we have previously defined the amino acid residues within a SLiM required for substrate binding to B55α. These residues direct dephosphorylation of a proximal phosphosite using p107 as a model substrate. We also shown that the SLiM is conserved in the unrelated substrate TAU and is required for its dephosphorylation. This allowed us to define an initial consensus B55 substrate SLiM, ‘p[ST]-P-x(4,10)-[RK]-V-x-x-[VI]-R’^14^. In this report, we have used a proteome-wide search tool, ScanProsite^24, 25^, to identify potential novel PP2A/B55 substrates with a degenerate version of this SLiM (p[ST]-P-x(4,10)-[RK]-[VIL]-x-x-[VIL]-[RK]) that includes potential conservative amino acid variants (Fig. 1A). This search yielded 275 proteins (1.3% of the proteome). We then filtered the list of potential substrate candidates by determining which of these proteins have been detected by B55α pull-down proteomics^10^, identified in phosphoproteomic analyses where B55α activity was inhibited in lysates^1^, or in phosphoproteomic datasets of proteins dephosphorylated following doxycycline-inducible expression of FLAG-B55α in HEK293 cells (Suppl. Table 1).

**Fig. 1.**
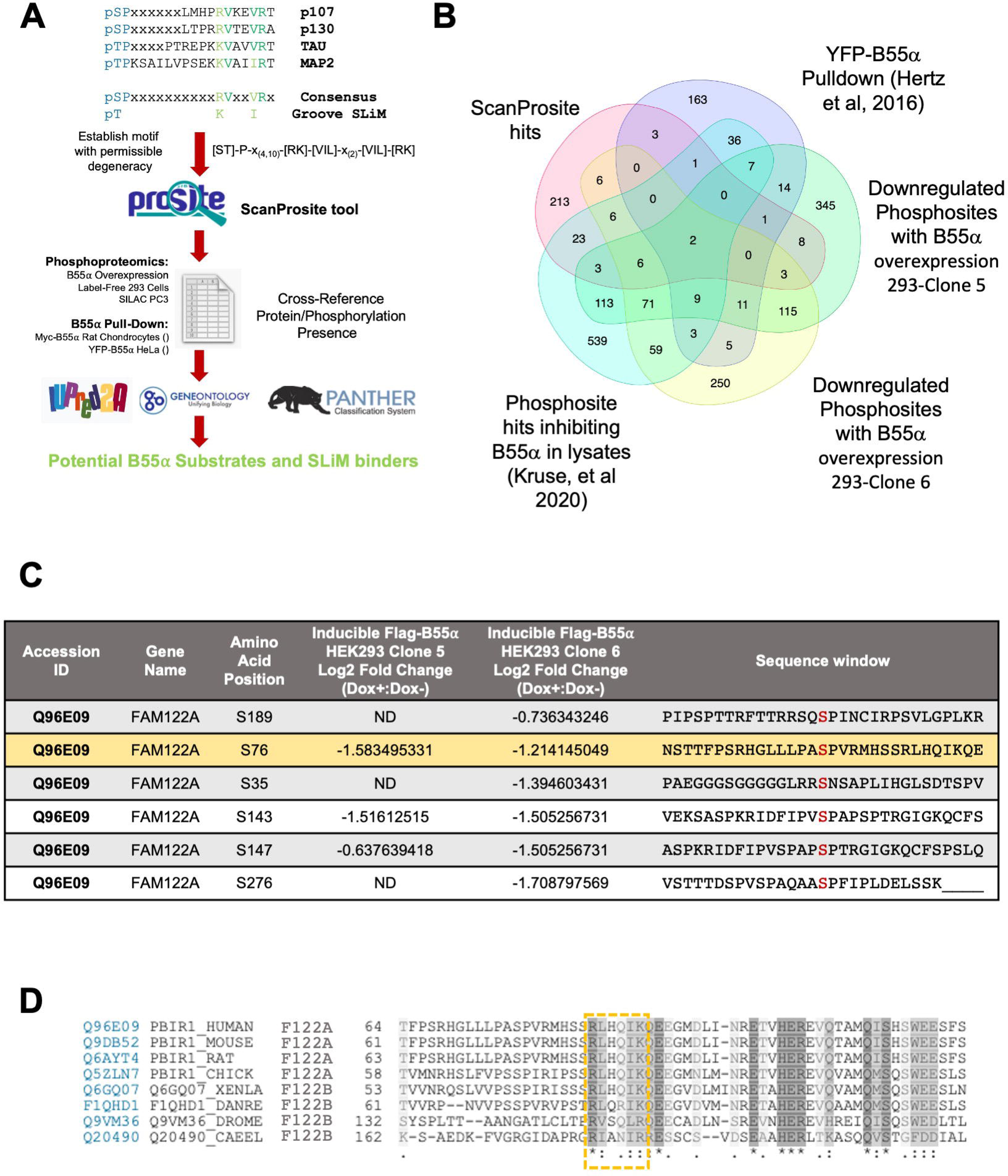
Strategy for identification of FAM122A as a SLiM containing protein and its conservation in bilateral animals. **A.** Steps for identification of PP2A/B55α substrates and other binders that use the SLiM. **B.** Venn diagram showing common hits among the ScanProsite search and proteins in the indicated datasets. **C.** FAM122A phosphosites downregulated upon doxycycline induction of B55α expression in HEK293-iB55α clones 5 and 6. **D.** The SLiM is conserved in bilateral animals: worms, flies, amphibians, birds, and mammals. The SLiM is boxed.

A recurring ‘hit’ among the datasets was FAM122A (Fig. 1B), also known as PABIR1 or PPP2R1A-PPP2R2A-interacting phosphatase regulator 1, a highly conserved protein of predicted disorder and a proposed inhibitor of the PP2A/B55α holoenzyme^22^. FAM122A exhibits a potential SLiM sequence, ***RL****HQ**IK***. FAM122A was identified in at least two independent B55α pulldown datasets^10, 17^, and exhibits several p-SP sites that are downregulated following upregulation of B55α in 293 and PC3 cells (Fig. 1C) including a proximal phospho-SP amino terminal from the potential SLiM sequence. Of note, a B55α pulldown dataset in Rat Chondrosarcoma cells, which we had previously reported, showed that FAM122A is the most abundant protein in B55α complexes other than the holoenzyme subunits^17^. Ontology analysis revealed that *FAM122A* originated by duplication of an ancestral *FAM122B* detected in *bilateria* present in worms, flies, frogs, and fish (Suppl. Fig 1A). Amino acid sequence conservation shows that the SLiM identified in FAM122A is conserved even in distant worms and flies (Fig. 1D; Suppl. Fig 1B), which also express B55/PP2A holoenzyme subunits and in the two FAM122A paralogs, FAM122B and FAM122C. Altogether, these data suggest that FAM122A is a potential SLiM-dependent inhibitor.

### FAM122A is a SLiM-dependent interaction partner of B55α

Since FAM122A has been reported to be a *bona fide* inhibitor of PP2A/B55α^22^, we sought to determine if its interaction with B55α is dependent on functional SLiM sequences via immunoprecipitation of cotransfected Myc-B55α and FLAG-FAM122A tagged mutants in transfected HEK293T cells. Mutation of murine mFLAG-FAM122A R81/L82 or I85/K86 SLiM residues to Ala abolished binding to Myc-B55α in reciprocal immunoprecipitations (Fig. 2A). In contrast, an S73A mutation, abolishing a phosphosite amino terminal to the SLiM, had no effect. As expected for a functional SLiM, the B55α D197K mutant, which does not bind B55α substrates^14^, also failed to bind FLAG-FAM122A (Fig. 2A). D197 is located in a deep groove on the surface of B55α, adjacent to the active site of the catalytic subunit. To rule out the possibility of the positively charged residues solely being responsible for the interaction, non-polar residues, Leu and Ile, were individually mutated to Ala in a GST-FAM122A deletion construct and subject to a GST-pulldown assay using HEK293T lysates. As shown in Fig. 2B, both L82 and I85 are critical for binding the holoenzyme *in vitro.* However, a short FAM122A_L70-K86_ peptide fused to GST containing the ‘***RL****HQ**IK’*** SLiM was not sufficient to bind the holoenzyme *in vitro* (Suppl. Fig. 2A), suggesting that additional residues contribute to binding.

**Fig. 2.**
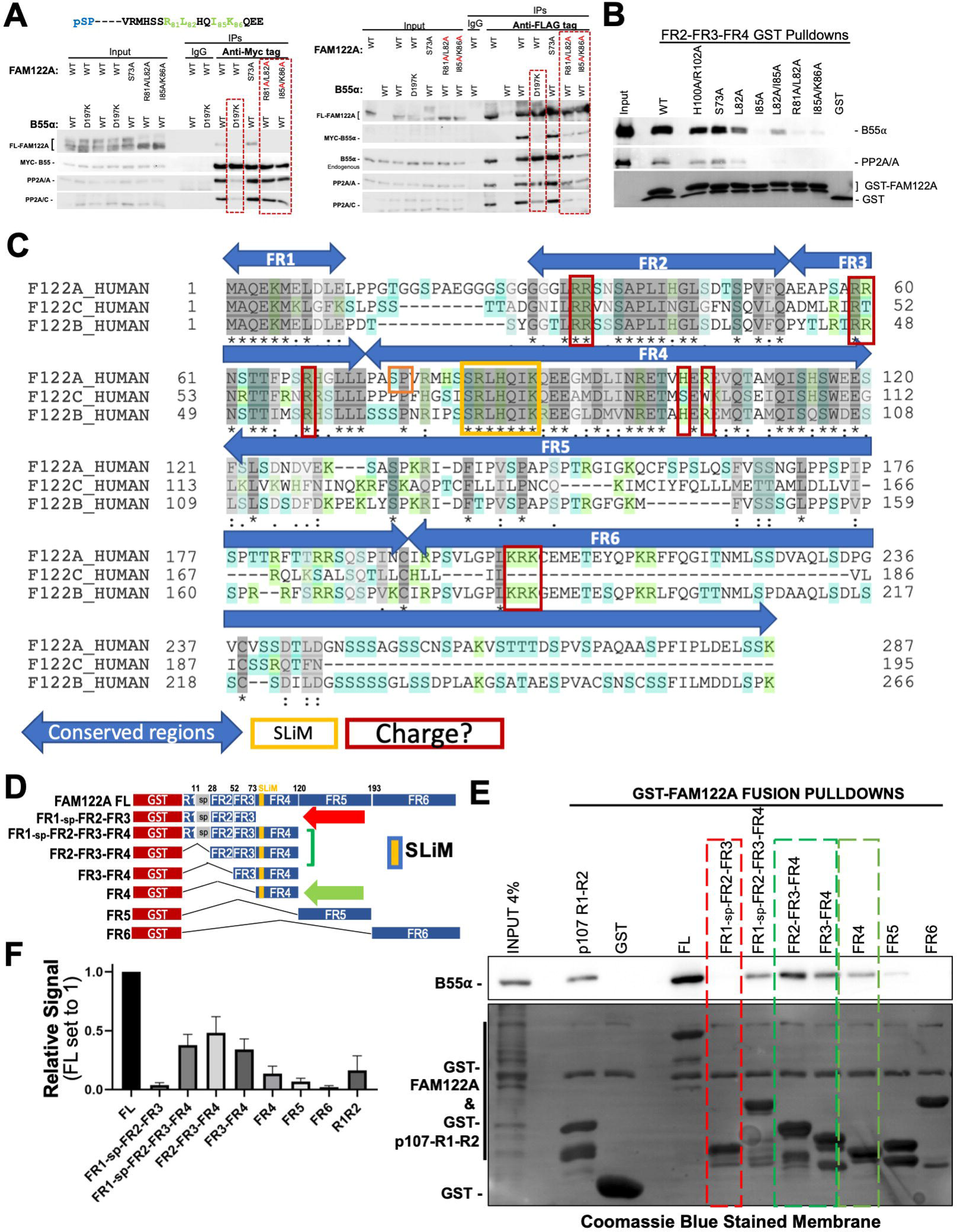
SLiM residues are required for FAM122A binding to B55α. **A.** HEK293T cells were cotransfected with FLAG-FAM122A and Myc-B55α WT and MTs. Anti-Myc and -FLAG IPs were analyzed by western blot. **B.** Glutathione beads loaded with WT and MT GST-FAM122A deletion constructs were incubated with purified PP2A/B55α and the pulldowns were analyzed by western blot. **C.** Human FAM122 family amino acid sequence alignment (FAM122A: Q96E09, FAM122C: Q6P4D5, and FAM122B: Q7Z309-3). Regions were selected based on conservation; the SLiM and residues that could mediate dynamic charge-charge interactions are marked. **D-F.** GST-FAM122A pulldown assays with the indicated constructs were done as in panel B and quantitated (F), n=4.

We next determined if other conserved regions in FAM122 family members (FR) were required or contributed to binding. FRs were selected by aligning the human FAM122A paralogs, FAM122B and FAM122C and denoting regions of conservation (Fig. 2C). Fig. 2D schematically details the collection of GST constructs used in pull down assays of HEK293T lysates. Conserved region FR4, which contains the SLiM, is sufficient for binding, while regions FR1, FR2, and FR3, fail to bind B55α, but FR2 and FR3 significantly increase the binding avidity of FR4 (Fig. 2D-F). As charge-charge dynamic interactions in PP2A/B56 substrates increase binding avidity to B56^26^, we mutated conserved positive residues in vertebrate FAM122A. Mutation of these conserved Arginine residues in FR2, FR3, and FR4 reduced binding to B55α (Suppl. Fig. 2B). Taken together, the interface of FAM122A:B55α binding is mediated via the SLiM and enhanced, at least in part by additional interactions of charged residues in FAM122A.

### *AlphaFold2* predicts that the hFAM122A SLiM folds as a short helix, followed by a longer C-terminal helix, and that R84 (corresponding to murine R81), contacts B55α D197

Despite previous computational metrics denoting disorder in FAM122A, we used *AlphaFold2*, an artificial intelligence (AI) program developed by Alphabet’s/Google’s DeepMind that performs predictions of protein structure^27^, to define any regions of human FAM122A that could fold. Remarkably, the SLiM is found within a predicted helix_R84-G93_ followed by a C-terminal helix_L95-S120_ (scoring confident to very high confident prediction, Fig. 3A) whereas the rest of FAM122A was predicted to be disordered. This finding prompted us to use ColabFold^28^, which implemented a version of *AlphaFold2* able to model heterodimers (“*AlphaFold2_advanced*”) with the original *AlphaFold2* program trained on single protein chains. This implementation, which became available in mid 2021 before the availability of AlphaFold-Multimer in late 2021, tricks *AlphaFold2* into modeling a complex by inserting 200 amino acids positions between the end of one protein sequence of interest and the beginning of another. Without the use of templates, *AlphaFold2_advanced* predicted the beta-propeller folding of B55α, which is very similar to the previously determined structure (PDB:3dw8)^13^. Four of five models placed the predicted short helix of FAM122A at the mouth of the B55α top groove (Fig. 3B, Suppl. Fig 3A). In the top model, the side chain of R84 (**R**LQHIK) forms a salt bridge (at 2.9 Å) with the side chain of B55α D197. L85 and I88 of FAM122A make hydrophobic contacts with B55α L225 (Fig. 3B-D); consistent with this, an L225A mutation of B55α reduces binding to FAM122A in cells (Fig. 3E).

**Fig. 3.**
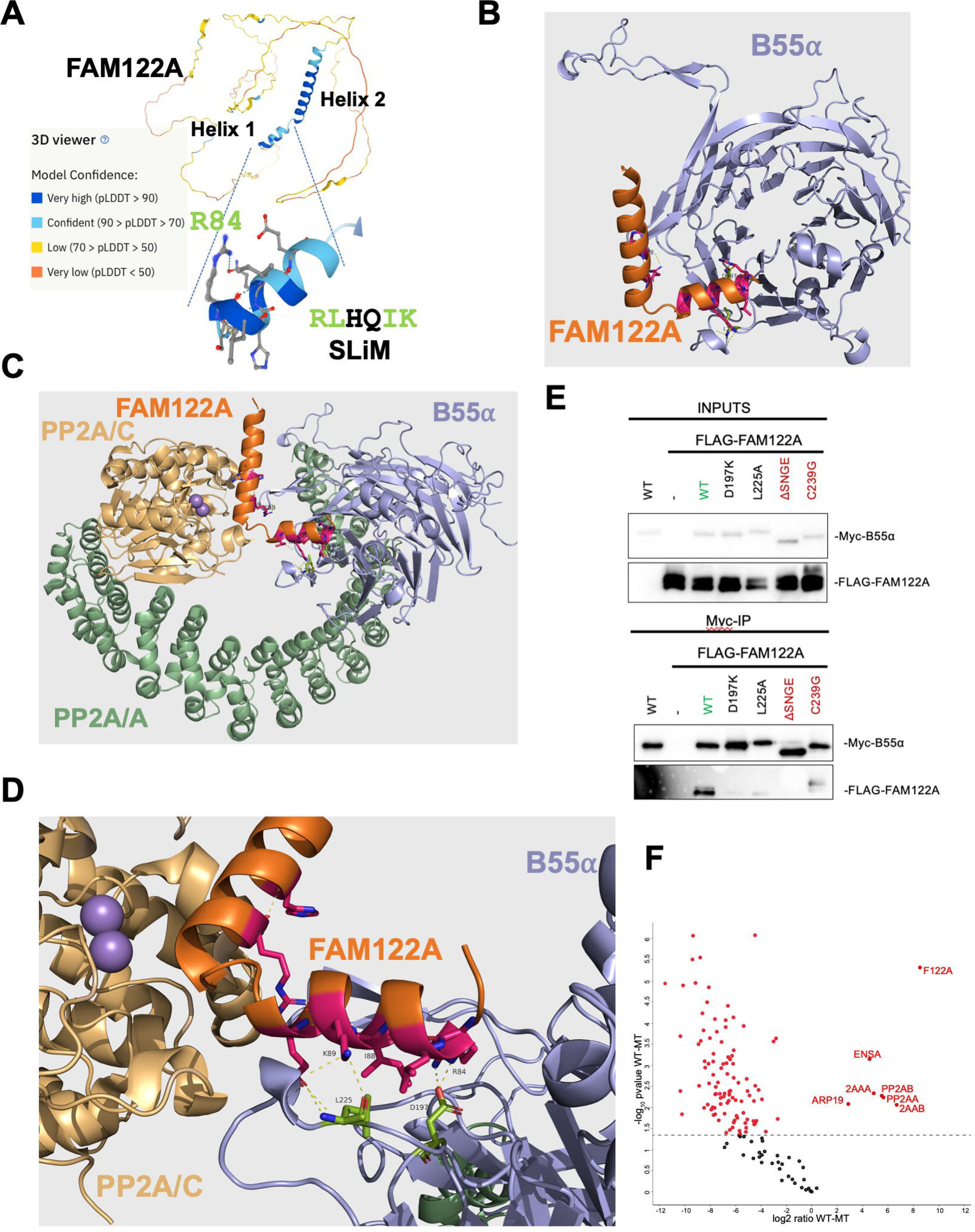
*AlphaFold2* predicts folding of a short helix in FAM122A containing the SLiM including R84 that makes contacts with D197 in B55α, with helix-2 blocking the active site. **A.** Original *AlphaFold2* predicts folding of two helices, with helix 1 containing the SLiM. **B.** *AlphaFold2_advanced* predicts that helix one binds at the mouth of a deep groove on the top of B55α that we have previously shown interacts with B55α substrates. **C.** The model was aligned to the structure of the holoenzyme (3DW8) by superposition of the B55α subunits in each structure. **D.** A closeup of the model aligned to the holoenzyme shows FAM122A R84 making contacts with D197 in B55α and L85-I88 with potential contacts to B55α L225. The position of helix-2 suggests a mechanism for blocking access to the active site and further stabilization of the enzyme/inhibitor complex. Residues are shown as numbered sticks: red for FAM122A, and lime for B55α. **E.** HEK293T co-transfections as in Fig. 2 with indicated mutants. **F.** Pulldown IP mass spec analysis of Myc-B55α-C239 vs Myc-B55α-wildtype. Relevant proteins are indicated.

In four of the five models predicted by *AlphaFold2_advanced*, the C-terminal long helix is predicted to be placed at a ∼90° angle to the SLiM-containing helix 1, suggesting that it could fill the space between B55α and the active site of PP2A/C. Therefore, we aligned the top predicted *AlphaFold2_advanced* B55α structure with the B55α subunit of the 3dw8 PP2A/B55α holoenzyme structure^13^. The superposition placed helix-2 in front of the active site (Fig. 3C). This strongly suggested that FAM122A would block substrate access to the PP2A/C active site. Moreover, residues within the long helix are predicted to be in close proximity with PP2A/C residues, suggesting additional contacts that could stabilize holoenzyme/inhibitor binding.

To gain insight into this possibility, we compared FLAG-FAM122A binding to wildtype (WT) Myc-B55α or a point mutant (C239G) that dramatically reduces B55α binding to the PP2A/A scaffold, rendering B55α monomeric in cells, yet capable of binding the p107 substrate (Suppl. Fig. 3B-D). This mutation reduced binding to FAM122A and resulted in a shift in mobility that could indicate changes in phosphorylation (Fig. 3E). In addition, immunoprecipitation from lysates of cells transfected with this B55α mutant vs WT followed by mass spec shows FAM122A as the most enriched holoenzyme binder (Fig. 3F, Suppl. Table 2). This indicates that FAM122A makes contacts with other subunits in the holoenzyme, consistent with the *AlphaFold2* model predicting contacts between the hFAM122A helix-2 (L_96_-S_120_) and PP2A/C. Altogether these data suggest that contacts to the holoenzyme are important for complex stability in agreement with the *AlphaFold2* model.

### FAM122A abrogates p107 binding to B55α *in vitro* and in cells

A previous report suggested that FAM122A inhibits PP2A/B55α by a mechanism involving degradation of the PP2A/C catalytic subunit^22^. However, we have not seen any effect in the catalytic subunit of the B55α holoenzyme when we co-express FAM122A in cells (Fig. 2A). In contrast, we have observed that under comparable concentrations of FAM122A and p107, FAM122A pulls down more B55α *in vitro* (Fig. 2E), suggesting comparatively higher affinity. In agreement with this, purified HA-FAM122A strongly inhibits B55α:p107 binding *in vitro* at FAM122A concentrations comparable to the PP2A/B55α complex (nM range) and with vast excess of p107_585-691_ (μM range) (Fig. 4A-B) in *in vitro* competition assays. Additionally, a dose dependent increase in transfected FLAG-FAM122A in HEK293T cells displaces FLAG-p107 from Myc-B55α (Fig. 4C). At comparable FLAG antibody signals, FAM122A abolishes p107 binding, which is consistent with FAM122A using its SLiM to block substrate binding and other conserved residues to bind with higher affinity than B55α substrates (among all PP2A/B55α substrates tested by us, p107 binds the tightest to B55α).

**Fig. 4.**
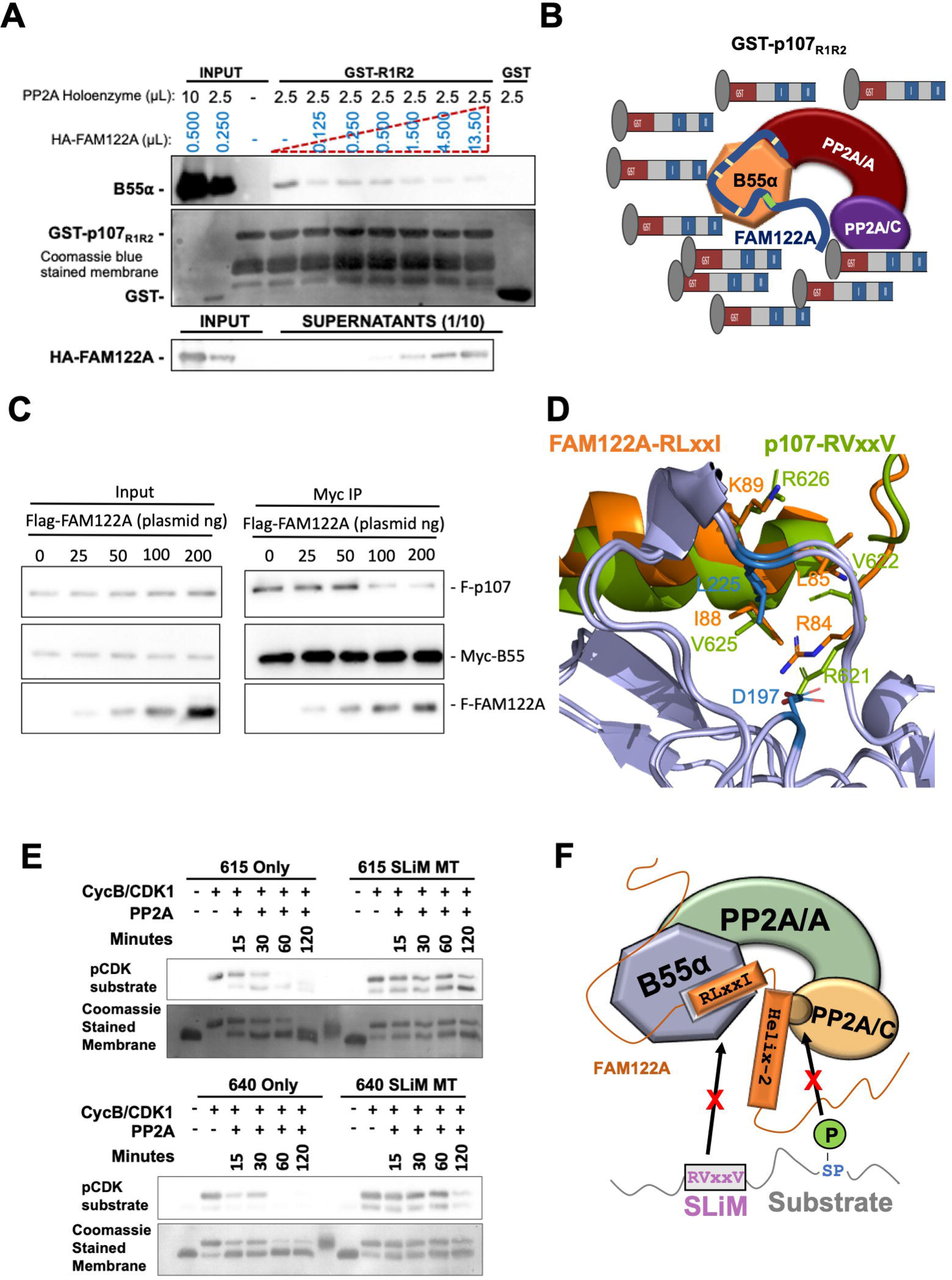
FAM122A effectively competes binding to the holoenzyme under conditions of high substrate excess. **A.** Purified PP2A/B55α holoenzymes were incubated with purified HA-FAM122A (expressed in *E coli*) starting with the same concentration as B55α in each sample up to a 100X. B55α:R1-R2 binding in GST-p107-R1R2 pull-downs was inhibited at the lowest FAM122A concentration. **B.** Schematic interpretation of A. **C.** Cotransfection of unvariable amounts of Myc-B55α and FLAG-p107 plasmids with increased amounts of FAM122A, as indicated. Anti-Myc immunoprecipitates were analyzed by western blot and relevant proteins are indicated. **D.** *AlphaFold-Multimer* model of superimposing a predicted helix containing the p107 with the predicted helix-1 of FAM122A. Conserved SLiM residues in both proteins superimpose and make contacts with D197 and L225. Residue numbering correspond to the human proteins. **E.** GST-R1R2 constructs created with a single phosphosite (S615 or S640). Arginine to Alanine mutations disrupted the SLiM motif in the indicated constructs. Western blot visualization of time dependent kinase-dephosphorylation assays of the R1R2 constructs. GST-R1R2 variant beads were phosphorylated with CDK1/cyclin B kinase. Washed phosphorylated beads were incubated with PP2A/B55α for the indicated times. **F.** Schematic representation of the substrate-competitive. mechanism of inhibition.

Further supporting a SLiM competition model, *AlphaFold2* applied to p107 alone also predicts a helix spanning the p107 ***R_621_V****KE**V**R* SLiM sequence, which prompted our use *AlphaFold-Multimer*^29^ to predict the interaction of p107 residues 612-648 with B55α. The original *AlphaFold-Multimer* was trained on multi-protein complexes in the Protein Data Bank deposited before April 2018. The model shows a helix formed from residues 620-633 which exactly superimposes the SLiM of p107 with that of FAM122A, with interaction of R621 with D197 and V625 with L225 (Fig. 4D). This binding model is compatible with the placement of p107 P-S640, located C-terminally with respect to the SLiM, at the PP2A/C active site. We therefore sought to determine if dephosphorylation of S640 was, like S615, dependent on the SLiM using p107-R1-R2 mutant variants that only contain a single SP-phosphosite. To make the phospho-S640 recognizable by an anti-p-SP antibody we mutated the residue preceding S640 to Met. Fig. 4E shows that S640 is dephosphorylated with kinetics comparable to S615, and that dephosphorylation of both phosphosites is dependent on the SLiM. Fig. 4F schematically summarizes the FAM122A competitive mechanism of inhibition.

### FAM122A SLiM folding as an α-helix is necessary for binding to B55α

As shown above, *Alphafold2* predicted that the SLiMs in both FAM122A and p107 are within a short helix. A representation of mFAM122A α-helix-1 is shown in Fig. 5A. We have previously shown that substitution of the two non-conserved residues within the p107 SLiM, K623 and E624, to Ala had no effect on binding to B55α^14^. These residues are not predicted to make contacts with B55α, and their substitution to Ala should not disrupt the α-helix. Consistently, mutation of mFAM122A ***RL****HQ**IK*** to ***RL****AA**IK*** had no effect on FAM122A binding to B55α in transfected HEK293T cells (Fig. 5B). We then examined if substitutions of these residues to Pro or Gly, which are typically not found in α-helices, would predict disruption of the short α-helix containing the SLiM in FAM122A. The top five *Alphafold2* models of FAM122A alone predicted that a FAM122A SLiM variant with Pro substitutions (***RL****PP**IK***) would not fold as a complete α-helix (Suppl. Fig. 5A). A FAM122A SLiM variant with Gly substitutions (***RL****GG**IK***), was predicted to still form an α-helix (Suppl. Fig. 5A). However, *AlphaFold2*’s predicted accuracy values (“pLDDT”) for the Gly residues were much lower (average value = 59.5 on a scale of 0-100) than for the WT residues (82.3). Residues with a value below 60 are considered unreliably placed. These results are not entirely surprising, since Gly residues are tolerated in α-helices but raise the entropy of the unfolded state, thus stabilizing the unfolded state (breaking the helix) relative to the α-helix form^30^. As predicted, FLAG-FAM122A variants with α-helix breaker residues, Pro and Gly, failed to bind Myc-B55α in anti-FLAG immunoprecipitations of lysates of HEK293T cells (Fig. 5B). These results strongly support the *Alphafold2* helical model, which spatially positions the SLiM residues forming a face that mediates contacts with B55α D197 and L225 (Fig. 4D and 5A).

**Fig. 5.**
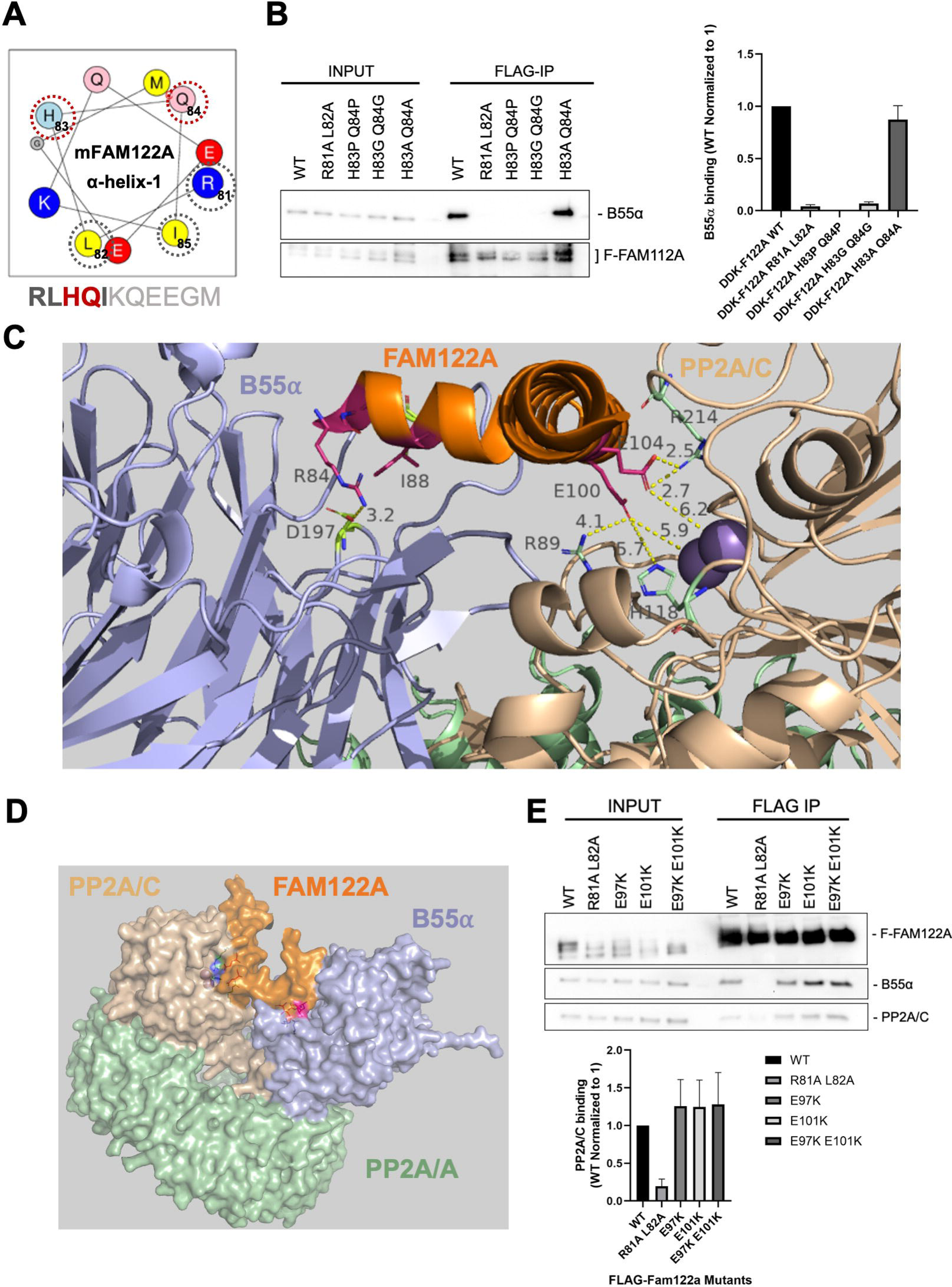
The FAM122A SLiM sits within α-helix-1 and contacts B55α, while α-helix-2 contacts the active site. **A.** FAM122A α-helix-1 representation using the *HeliQuest* tool. The amino acids that contact B55α are marked by a grey dashed circle. The variable residues mutated to Pro or Gly to break the helix are marked by a red-dashed circle. **B.** HEK293T cells were transfected with FLAG-FAM122A WT and helix tolerant and breaker MTs. Anti-FLAG IPs were analyzed for binding to endogenous B55α by western blot and relative binding was quantitated (n=4). **C.** *Alphafold-Multimer* v2.3 model closeup of FAM122A and the three subunits of the PP2A/B55α holoenzyme. Distances for the predicted contacts between residues in FAM122A with B55α and PP2A/C are indicated. The distance of E100 and E104 to the Mn^++^ metals is also indicated. **D.** *Alphafold-Multime*r v2.3 Surface model of the PP2A/B55α holoenzyme bound to FAM122A_76-122_. **E.** HEK293T cells were transfected with FLAG-FAM122A WT and E97K and E101K charge reversal MTs. Anti-FLAG IPs were analyzed for binding to endogenous B55α by western blot and relative binding was quantitated.

### AlphaFold-Multimer v2.3 model of the FAM122A/Holoenzyme tetrameric complex: FAM122A E100 and E104 within α-helix-2 serve as a flapper stopper at the active site of PP2A/C

The *AlphaFold2_advanced* model of FAM122A/B55α complex aligned to PP2A/B55α holoenzyme structure (3dw8) (Fig. 3C and D) suggests that hFAM122A residues E100 and E104 (corresponding to mFAM122A E97 and E101) occupy the active site of PP2A/C. However, FAM122A binding to the holoenzyme is likely to change the curvature of the PP2A/A scaffold, which is known to change depending on binding of distinct B subunits and SV40^13, 14^. We had attempted to model the full complex of FAM122A and the trimeric holoenzyme with *AlphaFold-Multimer* versions 2.1 (released November 2021) and 2.2 (released March 2022) but neither program placed FAM122A onto the holoenzyme (not shown). However, in December 2022, DeepMind released *AlphaFold-Multimer* v2.3, which was trained on a much larger set of protein complexes, along with other improvements in the deep-learning model (https://github.com/deepmind/alphafold/releases). This program successfully built a model of the tetrameric complex of FAM122A with the holoenzyme (Fig. 5C, Suppl. Fig. 5B), consistent with our earlier model, created by superposing the *AlphaFold2*-generated model of B55α/FAM122A with the crystal structure of B55α+PP2A/C+PP2A/A (Fig. 3C). The predicted accuracy metric (iPTM) was 0.804, a value which indicates that the model is likely to be very reliable^31^. The *AlphaFold-Multimer* 2.3 model is similar but not identical to the earlier model (Fig 3C) and likely models the precise interactions of FAM122A with PP2A/C more accurately, since both proteins are present in the full prediction process. Fig. 5C shows that hFAM122A E100 and E104 are predicted to make contacts with the manganese ions in the active site and the invariably conserved residues in PP2A/C that coordinate substrate phosphate (R89, H118, and R214)^9^. A complete surface model of the PP2A/B55α holoenzyme bound to FAM122A predicts that FAM122A fills the space between the active site of PP2AC and B55α with contacts to both proteins (Fig. 5D). Supp. 5B (right panel) shows that some residues C-terminal to the alpha helix-2 make contact with one of the beta sheets of the B55α beta propeller. Residues 155-159 of FAM122A form a beta strand (magenta), interacting with residues 163-167 of B55α (cyan). These residues’ positions are strongly predicted by AlphaFold-Multimer but remain to be tested experimentally for their functional effect. The model is available at https://doi.org/10.5281/zenodo.7739038.

With this new model in hand, we made mFAM122A variants with charge substitution of E97 and E101 (equivalent to E100 and E104 in hFAM122A) singly and in combination. As expected, FLAG-FAM122A wildtype, but not the SLiM mutant (R81A-L82A) immunoprecipitated B55α. In contrast, the E97K, E101K, and E97K-E101K FLAG-FAM122A variants bound B55α and PP2A/C similarly to WT FAM122A (Fig. 5E). This suggests that the electrostatic interactions mediated by these two Glu residues are important to hold the α-helix 2 in a position that blocks access to the active site, but insufficient to bind the PP2A/B55α holoenzyme in the absence of SLiM dependent interactions with B55α.

### FAM122A promotes proliferation

We next investigated the functional role of FAM122A in the cell cycle using CRISPR knockouts. FAM122A knockout in HEK293 cells dramatically reduces colony formation and proliferation (Fig. 6A-C). Inhibition of proliferation was also observed in T98G and U-2 OS cells using 2 different sgRNAs (Fig. 6C, Suppl. Fig 6A). Reconstitution of GFP-FAM122A in HEK293-FAM122A-KO cells partially rescued the proliferation defect in a SLiM dependent manner (Fig. 6D, Suppl. Fig 6B), indicating that the defects on FAM122A are at least partially dependent on PP2A/B55α inactivation. This is further supported as FAM122A knockout in HEK293 cells results in increased phosphatase activity relative to wildtype control in dephosphorylation of substrates phosphorylated on S/T-P sites (Suppl. Fig 6C).

**Fig. 6.**
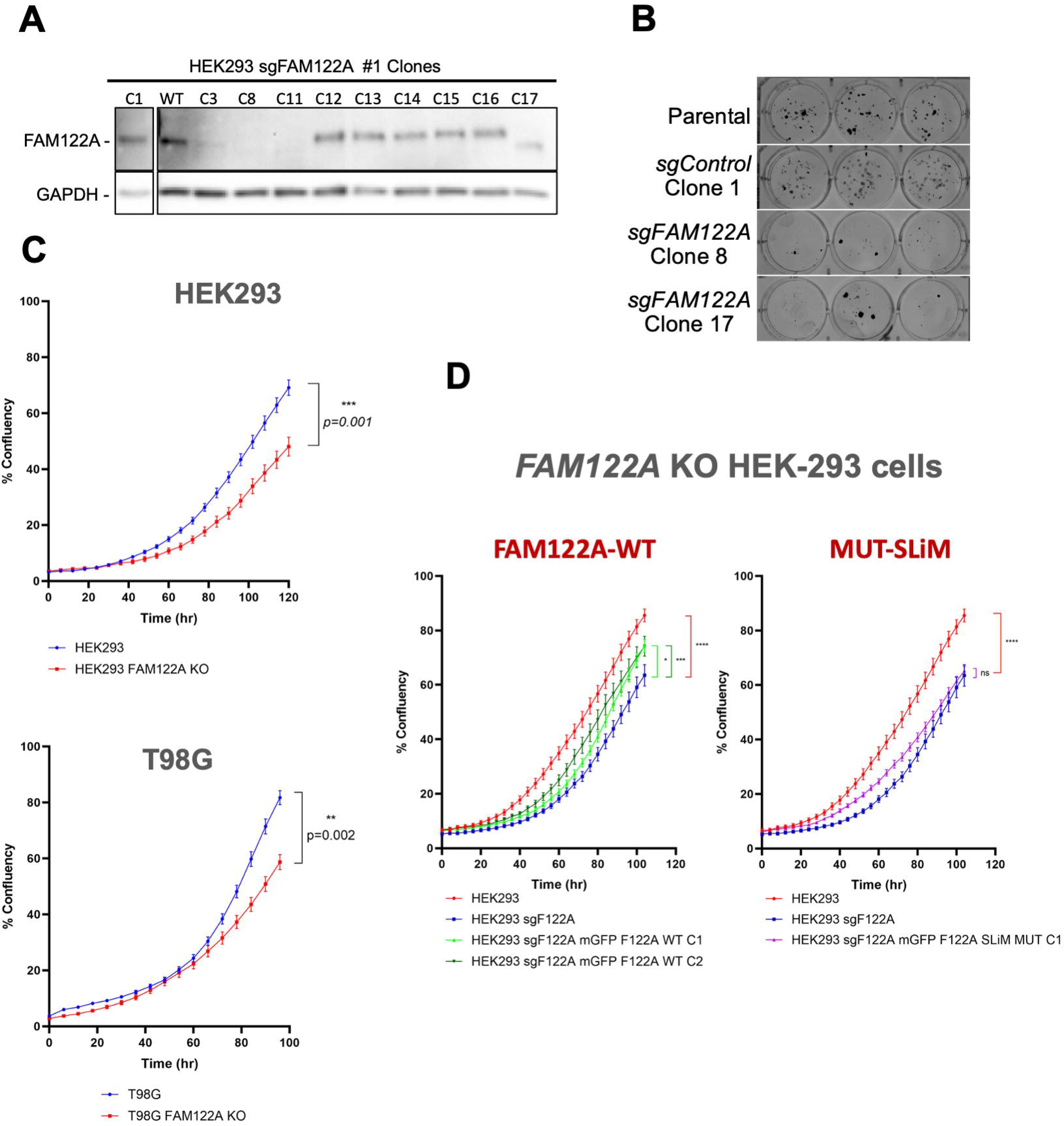
Ablation of FAM122A results in reduced proliferation that is rescued by re-expression of FAM122A. **A.** Western blot analysis of FAM122A CRISPR HEK293 clones (the C8 KO and C17 truncation were selected for further analysis). **B.** Colony formation assays of control (parental and C1) and KOs (C8 and C17). **C.** Ablation of FAM122A in HEK293, T98G cells inhibits proliferation. Cells were seeded in triplicate and percent confluence was imaged with an *Incucyte SX5*. **D.** A cassette directing the expression of FAM122A WT and SLiM-MT was stably introduced in HEK293 FAM122A KO cells. Expression of FAM122A wildtype but not SLiM mutant rescued the proliferation defect. HEK293 vs KO p value <0.0001 (****), HEK293 WT Clone 1 p value = 0.0436 (*), HEK293 WT Clone 2 p value = 0.0002 (***), HEK293 SLiM MUT Clone 1 p value =0.2640 (ns).

### FAM122A is required for timely cell cycle entry and progression through the G1 phase of the cell cycle following mitogen stimulation

Because PP2A/B55α opposes CDK inactivation of the pocket proteins, pRB, p107, and p130^17, 18, 19, 32^, and we have shown that PP2A/B55α directly dephosphorylates p107 *in vitro* and in cells^14^, we determined if knockout of FAM122A had any effects in cell cycle re-entry and progression through the cell cycle of T98G cells in response to serum stimulation. Parental and FAM122A KO T98G cells were serum starved for 3 days and then stimulated with serum. Cells were collected at the indicated points for DNA content FACS cell cycle and western blot analysis. FACS analysis shows a >4 h delay in the cells progressing from G0 to mitosis (Fig. 7A, nocodazole was added 16 h post-release to prevent cells from entering the next cycle). Western blot analysis showed a delay in the upregulation and reduced peak levels of Cyclin D1 that are consistent with the delays in pRB phosphorylation, the expression of E2F-dependent gene products (p107, Cyclins E and A) and the degradation of p130, which is dependent on CDK2 activity (Fig. 7B). To observe the delay more clearly, we performed a serum starvation and re-stimulation experiment where cells were collected hourly from 16 to 20 h. By 20 h the KO cells had not reached the number of cells in S phase already seen in the parental cells by 16 h (Fig. 7C). Similar delays in cell cycle progression and the expression of these markers were observed in cells where FAM122A was knockdown by siRNA (Suppl. Fig. 7A and 7B).

**Fig. 7.**
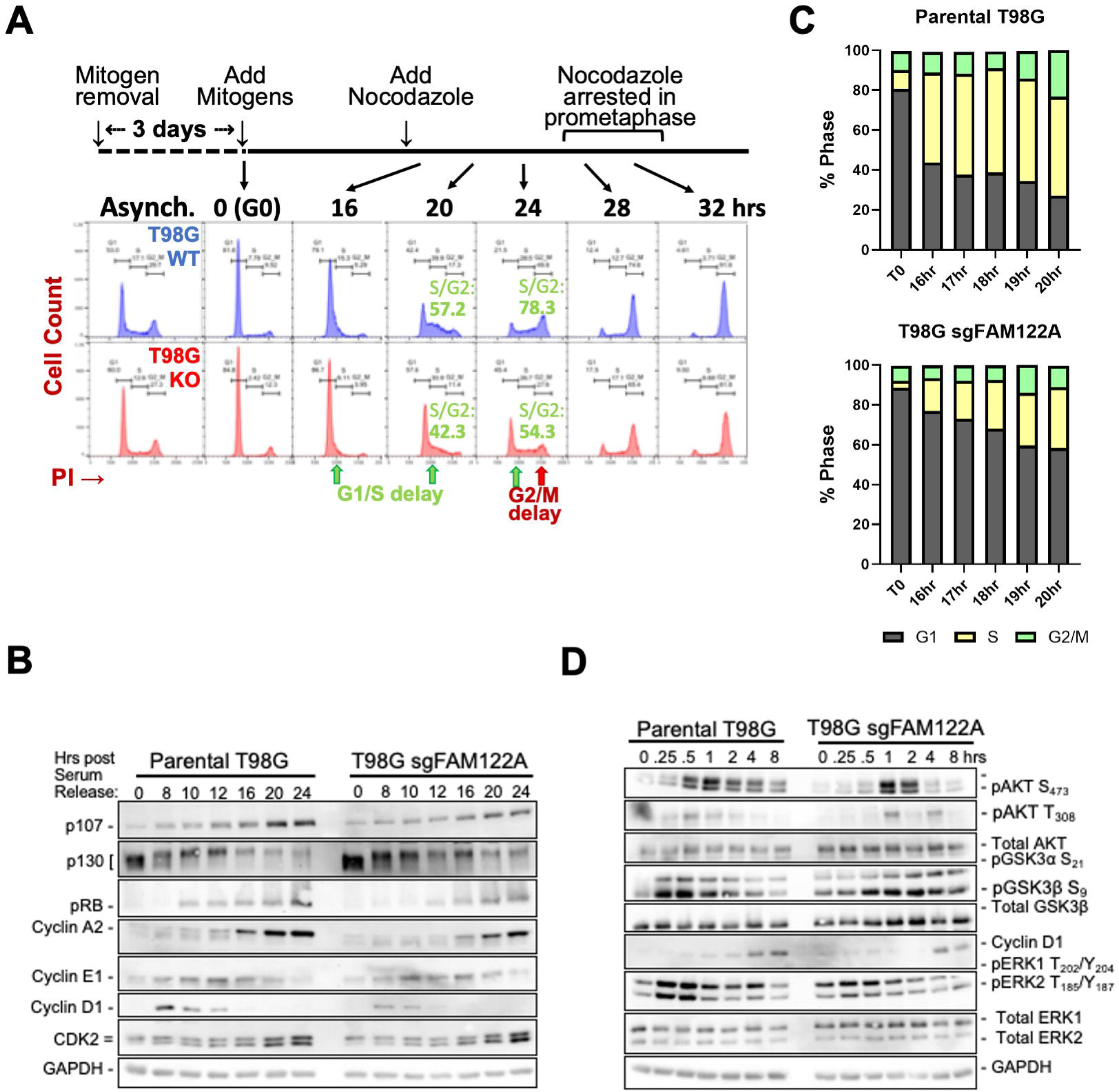
FAM122A is required for cell cycle entry and progression through the G1 phase of the cell cycle following mitogen stimulation. **A-B.** FAM122A-WT and -KO T98G cells were serum starved for 60 h and re-stimulated with DMEM supplemented with 10% FBS and collected at the indicated time points and analyzed by PI/FACS (A) and western blot analysis ( B). Nocodazole was added to prevent cells from progressing beyond mitosis. A >4 hr delay is observed by the time the cells reach mitosis, but the delay is already noticeable at the G1/S border. **B**. Delays in the expected modulation of pRB proteins and the expression of E2F-depedent gene products (Cyclins E and A and p107). The earliest defect is the limited expression of cyclin D1, whose expression is regulated by mitogens. **C.** A serum starvation-restimulation experiment focused on the G1/S transition (1 hr time points) that shows a delay in cells entering in S-phase. **D.** The effects of FAM122A KO in serum starved and restimulated T98G during early signaling and early G1 markers.

Because the decreased expression of cyclin D1 could explain a delay on pRB inactivation and passage to the restriction point and cyclin D1 expression is controlled by early mitogenic signaling^33^, we performed a serum starvation and re-stimulation experiment collecting cells at short time points preceding expression of cyclin D1. Lysates of these cells revealed delays in the activation of ERKs and AKT, both upstream regulators of cycling D1. The magnitude of peak signaling was also clearly lower (Fig. 7D). ERKs control cyclin D1 transcription, while AKT controls the stability of Cyclin D1 by inactivating GSK3β, which is known to promote Cyclin D1 degradation^33^. Consistently, inactivation of GSK3β is delayed in FAM122A-KO cells (Fig. 7D). These results, together with the FAM122A-SLiM dependent defects in proliferation (Fig. 7), strongly indicate that FAM122A controls PP2A/B55α activities that attenuate early mitogenic signaling including direct dephosphorylation of AKT T308, and other direct/indirect effects that oppose ERK activation (Suppl. Fig 7C).

### FAM122A is required for checkpoint activation in response to replication stress (RS)

Our data together with that published by others supports that FAM122A is an abundant regulated inhibitor of PP2A/B55α^3^. FAM122A has been recently shown to be phosphorylated and inhibited by CHK1 in NSCLC cell lines promoting activation of PP2A/B55α, which stabilizes WEE1 leading to inhibition of CDK1^23^. Interestingly, ablation of FAM122A restores WEE1 expression and the basal levels of inactive CDK1, restores fork replication speed, and dramatically reduces RS and DNA damage, suggesting that maintaining downstream control of CDK1 through FAM122A is the main mechanism by which CHK1 ensures genomic stability in these cells^23^. However, our data are only partially consistent with this model. Analysis of cell cycle effects of ablation of FAM122A in HEK293 cells using BrdU incorporation and propidium staining assays shows a clear reduction in the incorporation of BrdU to DNA during S phase, without a decrease in the number of cells with PI staining corresponding to S phase cells (Fig. 8A). This suggests that cells are synthesizing DNA more slowly and that the decreased proliferation in these cells, which have a defective pRB pathway, could be the result of extension of S-phase length, rather than accumulation of cells in G1. This also suggests that elimination of FAM122A is sufficient to induce replication stress, which was not reported in NSCLC cell lines^23^.

**Fig. 8.**
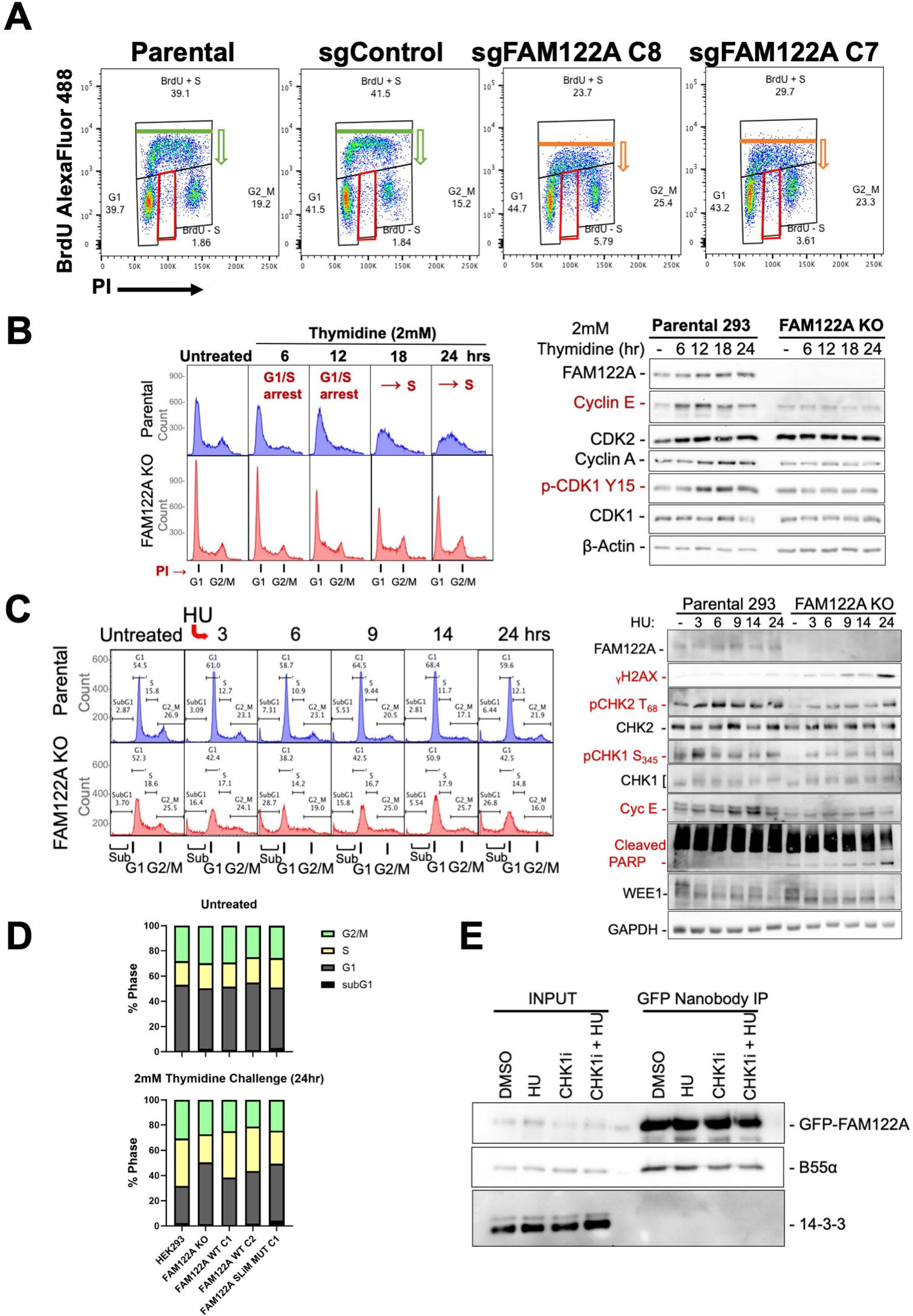
Ablation of FAM122A abrogates G1/S checkpoint activation in response to nucleotide depletion, which causes RS. **A.** Ablation of FAM122A KO in HEK293 cell results in a decrease in the overall incorporation of BrdU in cells in S phase and the appearance of a fraction of cells arrested in S-phase (BrdU negative, red boxes). **B.** HEK293 cells were treated with 2 mM thymidine for the indicated times and analyzed by PI flow and western blot using the indicated specific antibodies **C.** HEK293 cells were treated with 2 mM HU as indicated analyzed as in *B*. Relevant proteins are indicated. **D.** HEK293, FAM122A KO, and reconstituted WT expressing C1 and C2, and SLiM-MT expressing C1 cells were treated with 2mM thymidine and analyzed by PI flow as in panel B. **E.** GFP-FAM122A reconstituted HEK293 cells were treated with either DMSO, 2mM HU, 100nM CHKi, or in combination for 24hr and subject to GFP-nanobody pulldown for B55α and pan 14-3-3.

Consistent with this, while HEK293 are synchronized by either single or double thymidine blocks, FAM122A KO HEK293 cells fail to do so, which resulted in massive cell death without arrest of the cells in G1/S (Suppl. Fig. 8A). A thymidine challenge, which depletes endogenous nucleotides, resulted in time-dependent accumulation of cells at the G1/S transition and in S phase that correlated with accumulation of Cyclins E and A and inhibitory phosphorylation of CDK1 on Y15. By contrast, FAM122A-KO cells failed to arrest, and did not show changes in the expression of these markers (Fig. 8B-C). Similarly, nucleotide depletion by treatment with hydroxyurea (HU) resulted in effective G1/S and intra-S phase arrest with rapid activation of CHK2 (P-T68) and CHK1 (P-S345) and time-dependent accumulation of cyclin E in parental cells. This arrest was largely abrogated in FAM122A-KO cells, which exhibited rapid and time-dependent accumulation of γH2AX, sub-G1 DNA, and potent attenuation of both CHK2 and CHK1 activation (Fig. 8C). Of note, we did not observe changes in WEE1 expression, indicating that the G1 arrest mediated by FAM122A upon nucleotide deprivation is likely independent of WEE1. Altogether, these data strongly suggest that FAM122A controls PP2A/B55α-dependent steps upstream of these kinases. This is consistent with the ability of PP2A/B55α to negatively regulate ATM^21^, but likely involves other components in this pathway.

We next determined if the abrogation of the thymidine arrest checkpoint in HEK293 FAM122A KO cells could be rescued by reconstitution of FAM122A. Fig 8D shows that HEK293 FAM122A KO cells fail to arrest in S phase following 24 hr incubation with 2 mM thymidine. Reconstitution of WT FAM122A, but not FAM122A SLiM-mutant, restored the arrest in S phase. It has been recently reported that inhibition of CHK1 reduces phosphorylation of FAM122A presumably on S37, increases binding to B55α and reduces binding of 14-3-3 proteins, which results in FAM122A upregulation in the nuclear fraction of A549 lung cells^23^. FAM122A is also phosphorylated in cells at multiple sites that are consensus for the CDKs and ERK ^3^, indicating a potential regulation during the cell cycle. Therefore, we determined if FAM122A interaction with B55α is regulated during the cell cycle or in response to checkpoint activation. We did not observe changes in the levels of B55α bound to tagged FAM122A when comparing exponentially growing HEK293 and T98G cells, to cells arrested at the G1/S transition with thymidine, the G2/M transition with the CDK1 inhibitor (RO3306), and in pseudo-metaphase in cells arrested with nocodazole (Suppl. Fig. 8B-C). Also, we did not observe regulation of these complexes in T98G cells progressing through the cell cycle from quiescence or exiting mitosis and progressing through G1 from a nocodazole arrest (Suppl. Fig. 8D-E). These data suggest that the overall amount of FAM122A/B55α complex is constant under all these conditions and that regulation may depend on the colocalization of the complex and substrates and dynamic relative changes in FAM122A/substrate affinity for B55α.

We also determined if HU and/or inhibition of CHK1, which cause strong replication stress promote changes in the FAM122A/B55α interaction and binding to 14-3-3 proteins, which have been proposed to sequester FAM122A in response CHK1 phosphorylation^23^. We did not detect changes in the B55α/FAM122A interaction in GFP-FAM122A transfected cells (Fig. 8E). Surprisingly, we also did not detect formation of a FAM122A/14-3-3 protein complex upon induction of replication stress (Fig. 8E). Therefore, FAM122A is critical to establish checkpoints in response to replication stress that modulate CHK1 and CHK2 signaling,

Given the apparent lack of regulation of the FAM122A/B55α interaction as determined via immunoprecipitation, we determined the localization of FAM122A during the cell cycle. Transiently co-transfected RFP-FAM122A with EGFP-h-H2B in HEK293T cells revealed that FAM122A is largely, if not exclusively, localized in the nucleus in interphase, and it is released in the cytoplasm in prometaphase following nuclear envelop breakdown (NEB) (Fig 9A). Co-transfection of RFP-FAM122A, BFP-B55α and EGFP-h-H2B in 293T cells reveals that a fraction of B55α localizes in the nucleus and that both FAM122A and B55α are excluded from condensed chromosomes in mitosis (Fig. 9B). To demonstrate that B55α and FAM122A form a physical complex in the nucleus we used a Split-FAST approach^34^. B55α-N-FAST and FAM122A-C-FAST10 vectors were co-transfected in U2-OS cells and the FAM122A/B55α nuclear complex was detected upon incubation with the HMBR fluorogen (Fig 9C). Moreover, we used immunofluorescence to determine the localization of both FAM122A and B55α in the interphase and mitosis of U2-OS cells. As in HEK293T cells, FAM122A is nuclear in interphase, but it is not bound to chromatin in mitosis. A large fraction of B55α is also found in the interphase nucleus, and not bound to chromatin upon NEB. Of note, in late anaphase, both FAM122A and B55α appear to be recruited to chromatin (Suppl. Fig 9). Consistent with the nuclear localization of FAM122A, we identified a conserved nuclear localization signal (NLS, _201_RKK), that upon mutation (_201_RKK to AAK) prevented FAM122A from reaching the nucleus. Altogether these data show that FAM122A and B55α interact in the nucleus and that during mitosis, chromatin is devoid of FAM122A and B55α.

**Fig. 9.**
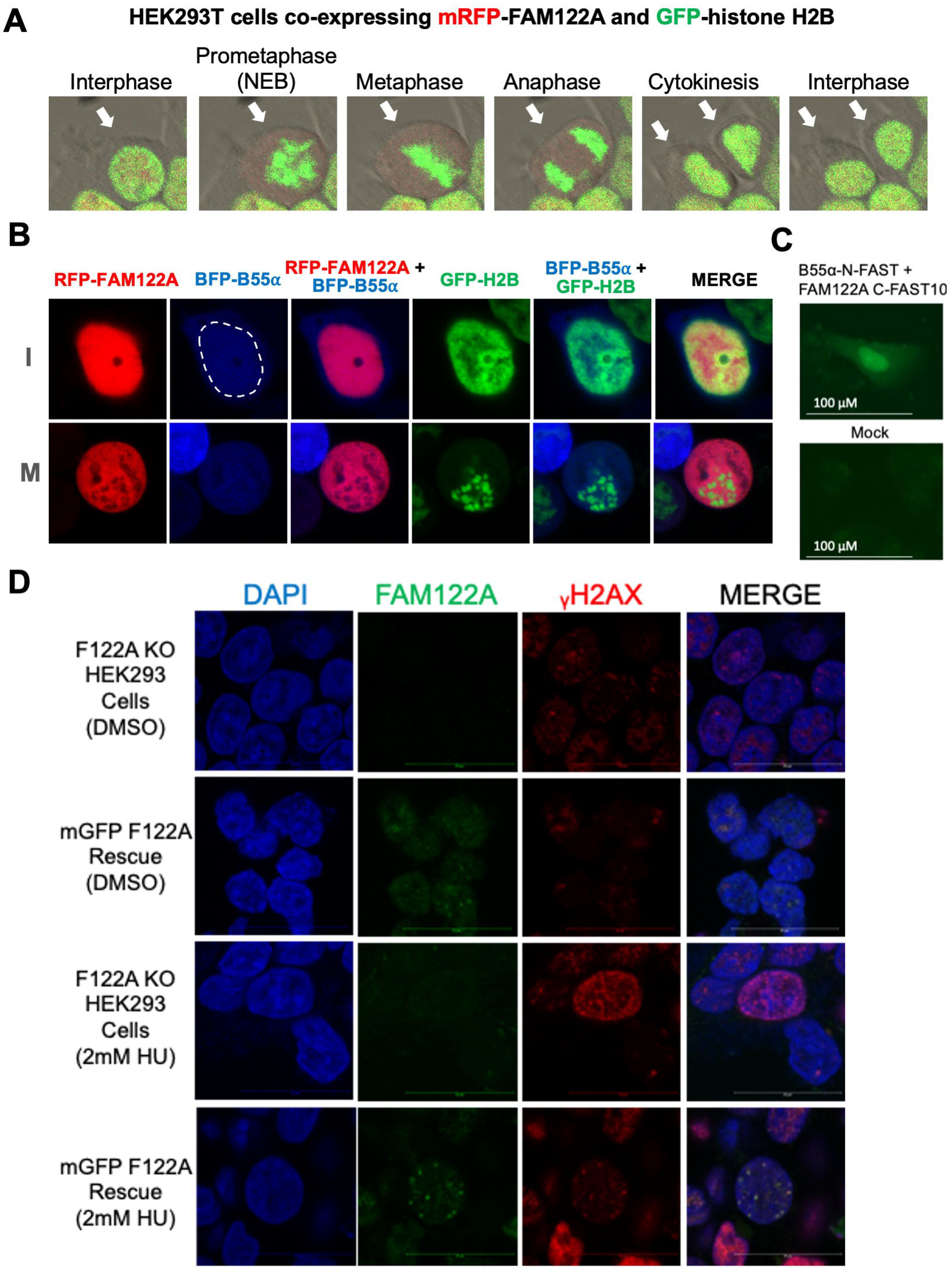
FAM122A is localized in the nucleus in interphase and to DNA damage *foci* in response to replication stress. **A.** mRFP-FAM122A wild-type was co-transfected with GFP-h-H2B into HEK293T cells and observed by time-lapse confocal microscopy (20x objective, 2.5x zoom factor). Arrows indicate the cell(s) in the cell cycle phase. **B.** mRFP-FAM122A, BFP-B55α and GFP-h-H2B expressing vectors were transfected into HEK293T cells to monitor localization. Close up of two cells in interphase (I) and in Mitosis (M) are shown. **C.** U2-OS cells were transfected B55α and FAM122A SplitFast constructs and colocalization was visualized by adding the HMBR fluorogen to the media 48 h later. **D.** FAM122A KO and reconstituted WT expressing C1 HEK293 cells were treated with 2mM HU for 24hr, stripped of cytoplasmic proteins and fixed for immunofluorescence imaging to observe localization of FAM122A bound to chromatin in the context of γH2AX foci on a confocal microscope using a 63x objective and 5x zoom factor.

Given the apparent lack of global regulation of the B55α/PP2A complex in response to replication stress, we tested the hypothesis that FAM122A localization to chromatin is regulated in response to replication stress. HEK293 FAM122A-KO with and without FAM122A reconstitution were treated with HU for 24 hours in glass chambers and soluble nuclear proteins were extracted with cytoskeletal buffer^35^. DMSO treated cells exhibit occasional γ-H2AX *foci* likely due to endogenous replication stress. Treatment with HU results in extensive DNA damage in the absence of FAM122A. Reconstitution of FAM122A in untreated cells results in clear localization of FAM122A at the chromatin and the larger foci colocalize with γ-H2AX. Treatment with HU resulted in strong recruitment of FAM122A to the sites of DNA damage (Fig 9.D). Therefore, these data suggest that FAM122A subnuclear localization is regulated and that the relative small fraction of FAM122A that is recruited to DNA-damage foci might be critical for establishment of the checkpoint.

## Discussion

Using a degenerate version of a consensus SLiM found in PP2A/B55α substrates p107 and TAU^14^, we have found that FAM122A, a previously identified inhibitor of PP2A/B55α, contains a functionally conserved SLiM sequence that is required for binding to B55α. FAM122A effectively competes with the model substrate p107 for binding to B55α *in vitro* and in cells. A previous report suggested that FAM122A binding to the PP2A/B55α holoenzyme results in downregulation of the catalytic subunit PP2A/C^22^, but we have not observed changes in PP2A/C binding to the holoenzyme upon FAM122A upregulation. The discovery and validation of the requirement of the SLiM residues in FAM122A allows a revision of our proposed SLiM to p[ST]-P-x(4,10)-[RK]_1_-[VL]_2_-x_3_-x_4_-[VI]_5_, but increased degeneracy at positions +2 and +5 is likely possible. As for the p107 SLiM, we find that a positively charged residue is not required at the final position for FAM122A binding to B55α. We have also shown here that the SLiM works to mediate dephosphorylation of sites both C-terminal and N-terminal from the SLiM, as both p107 S615 and S640, which flank the SLiM are dephosphorylated in a SLiM dependent manner.

We also show that the conserved SLiM is located in a region of FAM122A that is predicted by *AlphaFold2* to form an α-helix, where the **RL**xx**I**-SLiM residues R_84_ and L_85_-I_88_ mediate a salt bridge with B55α-D197 and hydrophobic contacts with B55α-L225, respectively. *AlphaFold-Multimer* also predicted an α-helix containing the p107 SLiM (**RV**xx**V**) interacting with the same residues on B55α, such that both the p107 and FAM122A α-helices superimpose. While SLiMs are sometimes thought to be extended structures, a few examples of helical SLiMs have been reported recently. For example, the C-terminus of the retinoblastoma protein pRB forms a 21 amino acid α-helix that has core residues that form a hydrophobic face^36^. If these residues are mutated to Ala, Cyclin D/CDK4 cannot phosphorylate pRB; however, substitution of other adjacent residues to Ala, which has high helical propensity^37^, had no effect. In contrast, disruption of α-helix by proline substitution, which prevents helix formation, impaired pRB dephosphorylation.

Our data-guided models predict that FAM122A simultaneously contacts both B55α and PP2A/C with dedicated α-helices 1 and 2. The SLiM residues in helix-1 are essential for binding. However, FAM122A E101 and E104, which are predicted to contact the manganese metal ions and the three invariable residues in all PPPs that coordinate substrate phosphate, are not essential for binding. This indicates that FAM122A E101 and E104 provide weak electrostatic interactions that allow helix-2 to occlude the active site. Additional contacts in the long IDRs upstream and downstream of the α-helices in FAM122A are likely to contribute to binding strength and may potentially be regulated by multiple phosphorylation sites that are known to be phosphorylated in cells in a cell-cycle-dependent manner^3^.

We have also noted that in the FAM122A-PP2A-holoenzyme complex, the curvature of the PP2A/A scaffold increases dramatically as compared to the PDB:3dw8 structure (Suppl. Fig 5C). We propose three potential mechanisms that are not mutually exclusive. First, the C-terminal residues in PP2A/C are present in the construct of the catalytic domain (sequence RRGEPHVTRRTPDYFL) in PDB:3dw8 but they are not ordered. *Alphafold2* is confident about the positions of those residues, which contact B55α (contacting residues 199-216 in B55α; average pLDDT score = 71.3). The C-terminal domain of PP2A/C is methylated and phosphorylation and methylation play a role in PP2A/B55α holoenzyme formation^6^, so it is tempting to speculate that methylation of L_309_ and/or phosphorylation of T_304_, Y_307_, may affect the interaction of B55α and the curvature of PP2A/A. Second, the presence of FAM122A bound to the holoenzyme may also directly affect PP2A/A curvature, bringing B55α closer to the active site. Third, the 3dw8 structure contains the cyanobacterial toxin microcystin in the active site of PP2A/C, which may increase the distance between PP2A/C and B55α, decreasing the curvature of the PP2A/A scaffold.

Ablation of FAM122A via CRISPR in multiple cell lines resulted in defects in proliferation and abrogation of the G1/S and intra-S checkpoint induced by replication stress. These defects were rescued by reconstitution of FAM122A in a SLiM dependent manner, demonstrating that FAM122A binding to B55α is essential for checkpoint activation. We also observed slower progression through the G0/G1 transition. Surprisingly, FAM122A interaction with B55α is largely constant during the cell cycle and upon mitotic checkpoint activation. Regulation may be restricted to modulation of FAM122A inhibitory function potentially through changes in FAM122A interactions regulated by posttranslational modifications within the IDRs. This was unexpected given recent work describing that CHK1 mediated phosphorylation of FAM122A results in its sequestration by 14-3-3 proteins and activation of PP2A/B55α that leads to WEE1 dephosphorylation and stabilization^23^. We have not observed 14-3-3 binding to FAM122A. Instead, we observe that replication stress induces formation of FAM122A *foci* at DNA damage sites marked by γH2AX. These data are also consistent with DepMap data that show inverse correlation in the effect of ablation of *PPP2R2A* with respect to *CHEK2 and TP53* in human cell lines (Suppl. Fig. 8F)^38^.

## Materials and Methods

### Cell culture and cell lines

All cell lines were obtained from ATCC and cultured in DMEM supplemented with 10% TET-Free FBS and 0.1% Penicillin-Streptomycin as described previously^18^ and tested for mycoplasma biannually or if there were any signs of stress. For lentivirus generation, HEK293T cells were transfected with pCMV-VSVG, pCMV-deltaR8.2, and expression constructs. Lentiviral supernatants were concentrated up to 100-fold using Takara Bio Lenti-X Concentrator. To overexpress FLAG-B55α in HEK293 cells, cells were transduced twice with pCW57.1 FLAG-B55α 24 hrs from one another and selected with puromycin and clonally isolated to determine expression levels among clones post doxycycline expression (48 hr). To genetically ablate *FAM122A*, HEK293, T98G, and U-2 OS cells were transduced with one of two lentiCRISPRv2-*sgFAM122A* (#1/#2) and selected with puromycin for clonal isolation. For Dox-inducible mRFP-FAM122A WT/Mutant cell line generation, T98G *sgFAM122A* #2 KO cells were transduced with pCW57.1-mRFP-*Fam122a* WT/MUT lentiviruses and bulk sorted according to mRFP fluorescence following doxycycline induction (48 hr). For mGFP-FAM122A WT/SLiM MUT HEK293 rescue cells, cells were transfected with pCMV6-AN-mGFP-*Fam122a* WT/SLiM MUT constructs with Lipofectamine 3000 and selected with G418 (500 µg/ml) followed by clonal isolation.

### Plasmids

Plasmids used or generated in this study are described in the Key resources table. pCMV6-*Fam122a*-Myc-DDK was purchased from Origene. pCMV6-AN-DDK-*Fam122a* was generated using SgfI/MluI restriction sites from the precision shuttle system (pCMV6-AN-DDK-Pol Iota-A kind gift from Roger Woodgate Addgene #131228). pCMV6-AN-mGFP and pCMV6-AN-mRFP plasmids were kind gifts from Richard Katz with Fam122a inserted using SgfI/MluI restriction sites. pGEX2T-*Fam122a*, pCMV6-AN-DDK-*Fam122a*, pCMV6-AN-mGFP-*Fam122a*, pCMV6-AN-RFP-*Fam122a*, pCW57.1-mRFP-*Fam122a*, and pGEX2T-FRs variant mutants were generated by site-directed mutagenesis using QuikChange II XL Site-directed mutagenesis (Agilent) with primers listed in the Appendix table and subsequently validated by Sanger sequencing. pGEX2T-FR deletion constructs were generated by PCR cloning BamHI/EcoRI restrictions sites, digestion, and ligation to pGEX2T. pCW57.1 FLAG-B55α and pCW57.1-mRFP-*Fam122a* were generated by LR clonase reactions (ThermoFisher Scientific) from the ORFs cloned in pENTR1A. (BamHI/EcoRI for mRFP-*Fam122a* and BglII/EcoRI for FLAG-B55α).

### Transfections

Transient transfections of Myc- and FLAG-containing constructs was achieved using calcium phosphate transfection or Lipofectamine 3000. In the case of calcium phosphate, 5 µg plasmid DNA was added dropwise to 2× HEPES-buffered saline (HBS) solution (280 mM NaCl, 50 mM HEPES, 1.5 mM Na2HPO4, pH 7.05) with bubbling, followed by addition to cells treated with 25 mM chloroquine after a 30 min incubation period. Alternatively, the transfection of stable cell lines was performed using Lipofectamine 3000 according to manufacturer’s recommendations.

### siRNA Transfection

siRNA for human *FAM122A* and *NC1* (negative control 1) were purchased from IDT. 10nM final concentration of *siFAM122A* was transfected in T98G cells between 50-60% confluency using Lipofectamine RNAiMax and incubating for 48hrs in serum free conditions.

### Phosphoproteomic Mass Spec Analysis

HEK293 iFLAG-B55α clones 5 (high expresser) and 6 (lower expresser) were induced with 1µg/mL of doxycycline for 48hr. To identify potential substrates and downstream targets of B55α in +/−Dox treated HEK293-iFLAG-B55α cells, global phosphoproteomics analysis was performed as previously described^39^ Protein identification was performed by searching MS/MS data against the Swiss-prot human protein database using andromeda 1.5.6.0 built in MaxQuant 1.6.1.0^39, 40, 41^. Statistical analysis was performed in the Perseus framework^42^.

### Affinity-Purification Mass Spec Analysis

Proteins from affinity-purified pull-downs were precipitated using either 20% trichloroacetic acid (TCA), washed twice with acetone (Burdick & Jackson, Muskegon, MI) or the single-pot, solid-phase enhanced method^43^. Proteins were digested overnight in 25 mM ammonium bicarbonate with trypsin (Promega) for mass spectrometric analysis. Peptides were analyzed on a Q-Exactive Plus quadrupole equipped with Easy-nLC 1000 (ThermoScientific) and nanospray source (ThermoScientific). Peptides were resuspended in 5% methanol/1% formic acid and analyzed as previously described. Raw data were searched using COMET (release version 2014.01) in high resolution mode^44^ against a target-decoy (reversed)^45^ version of the human proteome sequence database (UniProt; downloaded 2/2020, 40704 entries of forward and reverse protein sequences) with a precursor mass tolerance of +/-1 Da and a fragment ion mass tolerance of 0.02 Da, and requiring fully tryptic peptides (K, R; not preceding P) with up to three mis-cleavages. Static modifications included carbamidomethylcysteine and variable modifications included oxidized methionine. Searches were filtered using orthogonal measures including mass measurement accuracy (+/-3 ppm), Xcorr for charges from +2 through +4, and dCn targeting a < 1% FDR at the peptide level. Quantification of LC-MS/MS spectra was performed using MassChroQ (Valot et al., 2011) and the iBAQ method^46^. A total peptide count > 1 at least two quantifications in either the WT or MT sample were required for inclusion in the dataset. Proteins with a total peptide count > 1 were removed as non-specific binders. Missing values were imputed, and statistical analysis was performed in Perseus^42^.

### Cell cycle, Flow Cytometry, and Cell treatments

Parental and KO T98G cells were serum starved for 3 days prior to mitogenic stimulation with serum-containing media and collected at the indicated timepoints for flow cytometry and WB analysis. Collection for flow cytometric DNA content analysis was performed by collecting the cells followed by ice-cold ethanol fixation. Following 2 PBS washes, cells were stained with 1x propidium iodide solution (in 1% FBS in PBS with 1mg/mL RNase A) in the dark for 30 minutes. For cells expressing mGFP (TagGFP), cells were fixed in ice-cold methanol to preserve GFP fluorescence and processed as samples fixed with 70% ethanol.

BrdU incorporation assays in HEK293 cells were performed by the addition of 10µM BrdU to cells for a 45-minute pulse. Cells were fixed using ice-cold 70% ethanol and left overnight for permeabilization. Cells were washed with PBS twice followed by denaturing with 2N HCl for 20 min. Following washes and neutralization with 0.1M Na2B4O7 for 10 min, cells were incubated with anti-BrdU antibody (1:200) for 20 min. Subsequent washes were followed up by anti-rabbit Alexa Fluor 488 secondary (1:200) for 10 min. Following two additional washes, cells were stained with 1x propidium iodide solution as described above.

All flow cytometry experiments were carried out on a BD LSR-II and analyzed in FlowJo v10 (BD Biosciences). For nocodazole release, cells were initially treated with 10 nM nocodazole for 20-24hr followed by washout and collections at indicated timepoints. For RO3306 (CDK1i), cells were treated with 10µM for 24hr and either collected or released following PBS washout and collected at given time points for PI staining as described above. For Thymidine and HU Challenge/Release experiments cells were treated with 2mM Thymidine or HU for given time points (24hrs in the case of release) and collected or released with collections at the given timepoints post-release. For CHK1i (prexasertib) treatments cells were treated with 100nM for 24hr and collected.

### Clonogenic growth and proliferation analysis

Proliferation curves were established by seeding 1000, 2000, 3000, and 4000 cells per well with 4 wells per condition and the area phase contrast measured every 4 hours with an IncuCyte SX5. For clonogenic assays, cells were cultured for 11 days followed by fixation and stained with crystal violet.

### GST Pulldown Assays

GST-tagged constructs of interest were expressed in Escherichia coli bacteria and purified for use in pull-down assays. Briefly, 100 mL cultures of E. coli were treated with 0.25 mM isopropyl β-D-thiogalactoside (IPTG) for 3 hr to induce expression of GST-fusion proteins. Cells were then harvested by centrifugation and resuspended in NETN lysis buffer (20 mM Tris pH 8, 100 mM NaCl, 1 mM EDTA, 0.5% NP-40, 1 mM PMSF, 10 µg/mL leupeptin) prior to sonication at 30% amplitude for 10 cycles. Supernatants were collected and incubated with glutathione beads for purification, followed by NETN buffer washes for sample clean-up. For pull-down assays, purified GST-FAM122A, deletion, and mutant constructs were incubated with HEK293T lysates for 3 hr or overnight at 4°C, followed by washes (4×) with complete DIP lysis buffer (50 mM HEPES pH 7.2, 150 mM NaCl, 1 mM EDTA, 2.5 mM EGTA, 10% glycerol, 0.1% Tween-20, 1 µg/mL aprotinin, 1 µg/mL leupeptin, 1 µg/mL Pepstatin A, 1 mM DTT, 0.5 mM PMSF) and elution with 2× LSB. Samples were resolved by SDS-PAGE and probed using antibodies against proteins of interest.

### In vitro Kinase and Phosphatase Assays

GST-tagged p107 constructs loaded on beads were phosphorylated with recombinant cyclin B/CDK1 (Invitrogen) in 200 nM ATP KAS buffer (5 mM HEPES pH 7.2, 1 mM MgCl2, 0.5 mM MnCl2, 0.1 mM DTT). We incubated samples with shaking at 37 °C for 2 hr unless indicated. Reactions were stopped by adding 2× LSB and heated at 65 °C when used for PAGE or immunoblots. In vitro phosphorylated substrates for phosphatase assays were washed 3× in complete DIP buffer, followed by addition of indicated concentrations of purified PP2A/B55α. Reactions were incubated with shaking at 37°C for times indicated, followed by addition of 2× LSB and boiling at 65 °C for western blotting.

### Eluted Fam122a for Competition Assays

A PreScission protease site and 2x-HA tag were added to pGEX2T-Fam122a via restriction enzyme cloning. GST-protein purification was performed as described above. 2x-HA-Fam122a was eluted off GSH beads by PreScission protease cleavage in PreScission protease cleavage buffer (50 mM Tris-HCl, pH 7.0 (at 25 °C), 150 mM NaCl, 1 mM EDTA, 1 mM dithiothreitol) at 4 °C overnight. Following brief centrifugation, the eluate was collected and prepared to contain 25% glycerol and 1 mM DTT for –80 °C storage. Briefly, purified PP2A/B55α holoenzymes were preincubated with increasing nM concentrations of Fam122a for 30 min at 37°C to facilitate interaction. These preincubated PP2A/B55α complexes were then incubated with µM concentration GST-tagged p107 R1R2 constructs for 3 hr or overnight at 4°C, followed by washes (4x) with DIP lysis buffer and elution with 2× LSB. Proteins were resolved via SDS-PAGE and detected via western blotting using anti-B55α and GST antibodies.

### Immunoprecipitations

Whole-cell extracts (200–1000 µg) were incubated with Myc- or FLAG-conjugated beads (Sigma) for 3 hr or overnight at 4 °C. Input samples were collected prior to antibody-conjugated bead incubation, and supernatants were taken post-incubation following sample centrifugation. Beads were then washed (4×) with complete DIP lysis buffer and proteins were eluted in 2× LSB. Proteins were resolved by SDS-PAGE and probed using antibodies against proteins of interest.

### Protein Analyses

Cells were lysed with ice-cold lysis buffer (50 mM Tris-HCl (pH 7.4), 5 mM EDTA, 250 mM NaCl, 50 mM NaF, 0.1% Triton X-100, 0.1 mM Na_3_VO_4_, 2 mM PMSF, 10 μg/ml leupeptin, 4 μg/ml aprotinin, and 40 μg/ml Pepstatin A) or Complete DIP. Proteins were resolved by 8% or 10% polyacrylamide/SDS gel electrophoresis, and transferred to a polyvinylidene difluoride (PVDF) membrane (Immobilon-P, Millipore) in 10 mM CAPS/10% methanol buffer (pH 11). Bands were visualized by using SuperSignal West Pico PLUS Chemiluminescent Substrate (Thermoscientific) and imaged using an iBright CL1000 imaging system (ThermoFisher). Antibodies used are listed in the key resource table. Densitometric analysis of western blots was performed in Fiji (ImageJ)^47^.

### Graphs and Statistical Analysis

All experiments were performed in triplicate unless otherwise specified. Graphs depict calculated SEM from all replicates. Graphs were generated in Prism Graphpad 8. Statistical analyses were performed using Student’s t-test and represented as follows: *<0.05, **<0.01, ***<0.001, and ****<0.0001. Statistical analyses of proliferation curves comparing parental to

FAM122A knockout cell lines was conducted using a Wilcoxon matched-pairs signed rank test of the last 10 measurements. In the case of WT FAM122A and SLiM MUT rescues, a Friedman test was performed of the last 10 measurements comparing all conditions to FAM122A KO cells.

### Immunofluorescence Staining and Fluorescent Microscopy

Cells were seeded on iBidi 8 well glass bottom (Cat. 80827) µ-slide treated previously with Poly-D-Lysine (Cat. A389040). Cells were treated with hydroxyurea (2 mM for 24 hr or 4 mM for 6 hr), 100 nM Prexasertib for 24 hr or in combination for 24 hr. Cells were treated as previously described^48^ for whole cell staining or as described for chromatin bound staining^35^ using α-FAM122A (1:50), α-ᵧH2AX (1:200), and α-B55α (2G9) (1:50) antibodies. Anti-rabbit AlexaFluor 488 and Anti-mouse AlexaFluor 594 secondary antibodies were used at 1:100 dilution. All confocal microscopy was performed on a Leica TCS SP8 microscope using 63x oil immersion lens. Other imaging was performed using an EVOS cell imaging system.

### AlphaFold2 calculations

Some calculations were performed using the Jupyter notebooks provided by ColabFold^28^, which requires inputting the query sequence and choosing options for running the calculation using Google Colab GPUs. We used both the original *AlphaFold2* notebook and the *AlphaFold2_advanced* notebook

(https://colab.research.google.com/github/sokrypton/ColabFold/blob/main/beta/AlphaFold2_adv anced.ipynb). The latter was developed to model complexes of proteins before the availability of *AlphaFold-Multimer*. The notebook https://colab.research.google.com/github/sokrypton/ColabFold/blob/main/AlphaFold2.ipynb implements both the original AlphaFold2 trained on single-chain proteins and *AlphaFold-Multimer* for complexes. We ran *AlphaFold2* without templates on the ColabFold notebooks.

For calculations with the recently released *AlphaFold-Multimer 2.3* (December 2022), we downloaded code from the DeepMind github repository (https://github.com/deepmind/alphafold) for computations on a Linux workstation with a 24 Gbyte GPU. All version of AlphaFold2 run five separate models consisting of the weight matrices that convert input features (such as the query sequence and the multiple sequence alignment of homologues) into a predicted structures. The first two models use templates. Models 3, 4, and 5 do not. Structure predictions for the tetrameric complex of PP2A/A, B55α, PP2A/C, and FAM122A were performed with the use of templates (PDB70) in models 1 and 2. Each set of 5 models from *AlphaFold-Multimer* was run with five random seeds for a total of 25 models. Models were ranked as recommended in the AlphaFold-Multimer paper^29^, which is a linear combination of the pTM and iPTM scores (0.8*iPTM + 0.2*pTM), which combines the “interaction” predicted TM score (iPTM) and the whole structure predicted TM score (pTM). The top model of the tetrameric complex had a iPTM+pTM score of 0.804. This structure prediction came from *AlphaFold-Multimer* model4, and so did not use experimental structures as templates. Structures were optimized with AMBER (default in AlphaFold2) and visualized with PyMOL. The model is available in Supplemental Data File 1.

The AlphaFold-Multimer model of the tetrameric complex and a PyMOL session are available at https://doi.org/10.5281/zenodo.7739038.

## Supporting information

Key Resources Table

## Acknowledgements

This work was supported in part by the National Institutes of Health Grants R01 GM117437 and R03 CA216134-01, a WW Smith charitable Trust Award, a FCCC/TU Nodal award (XG), R35 GM122517 (RLD), and a Pre-Pilot Award from U54 CA221704 (JW, ZZ and HF) and funding from NCI CCSG grant P30 CA006927 (XG, JD and RLD) that supports the Molecular Modeling Facility at the FCCC. We also thank Guercie Guerrier for technical assistance.

---

**Suppl. Fig. 1.**
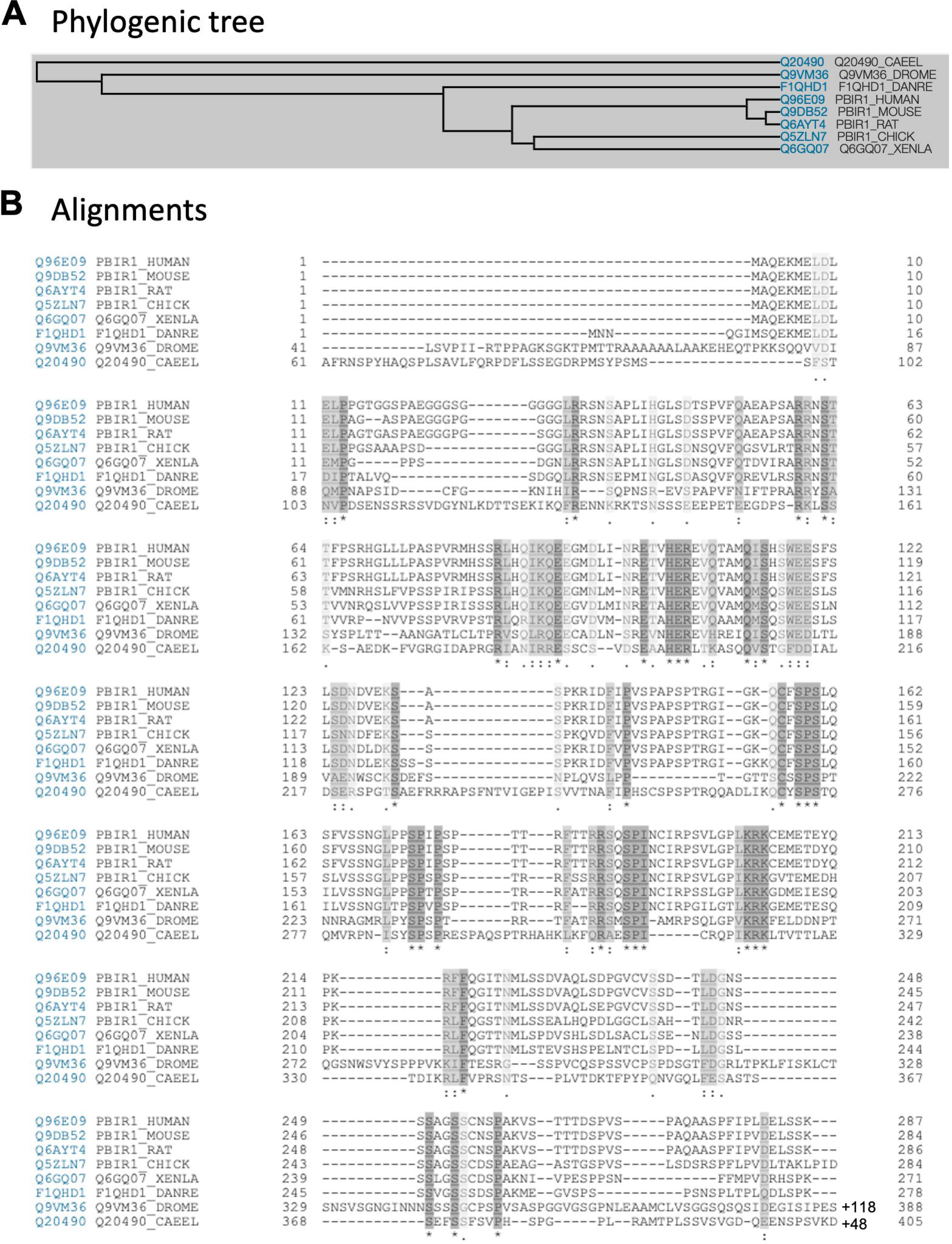
FAM122A is first found in *Bilateria* animals, and the SLiM is Fully conserved. **A.** Phylogenetic tree. **B.** Sequence alignment.

**Suppl. Fig. 2.**
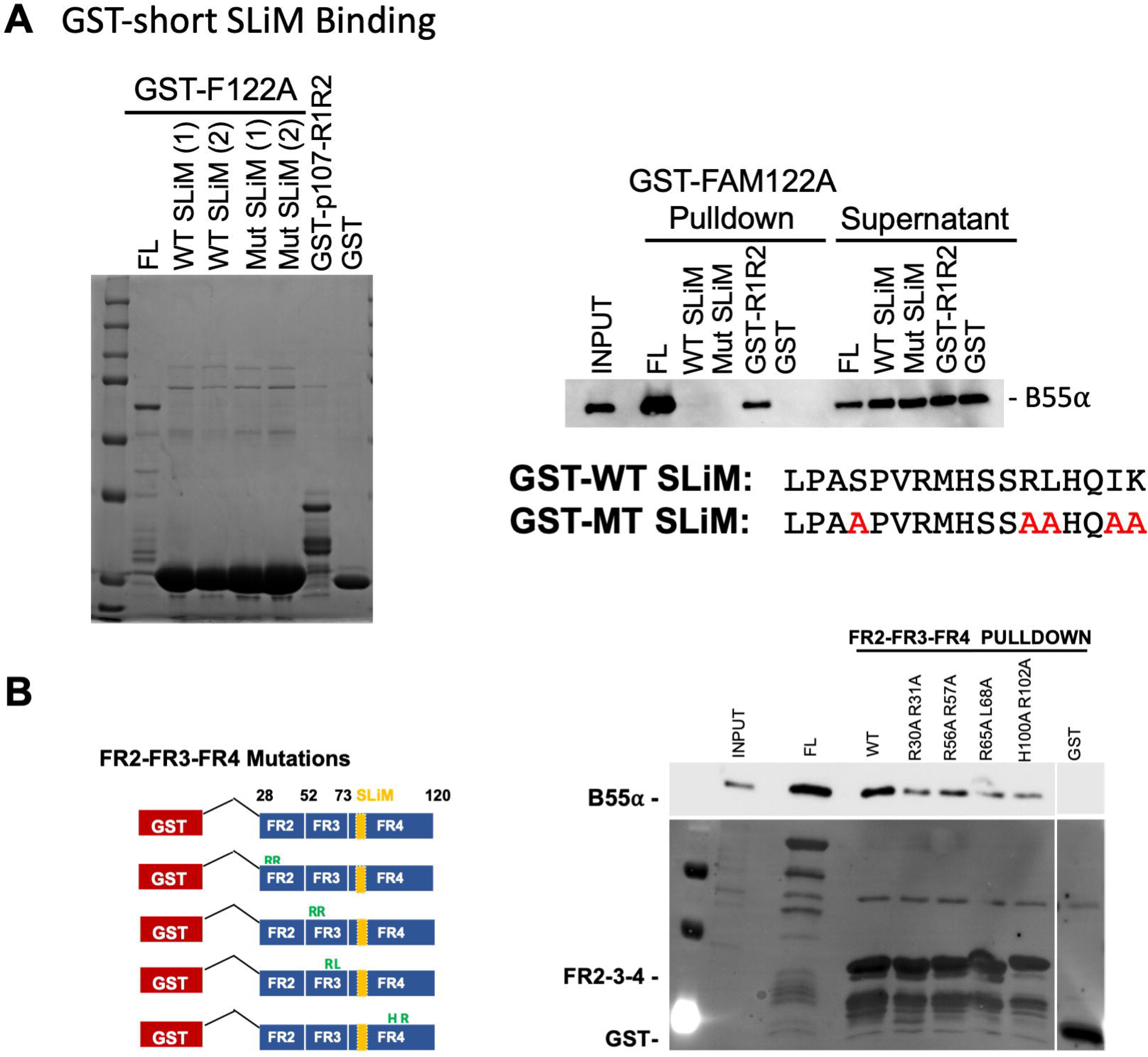
A. While the SLiM is essential for binding B55α, sequences beyond FR4 contribute contacts. **A.** A 17 residue peptide containing the SLiM is insufficient for binding B55α. **B.** GST-FAM122A pulldown assays with the indicated variant constructs eliminating positively charged amino acid pairs.

**Suppl. Fig. 3.**
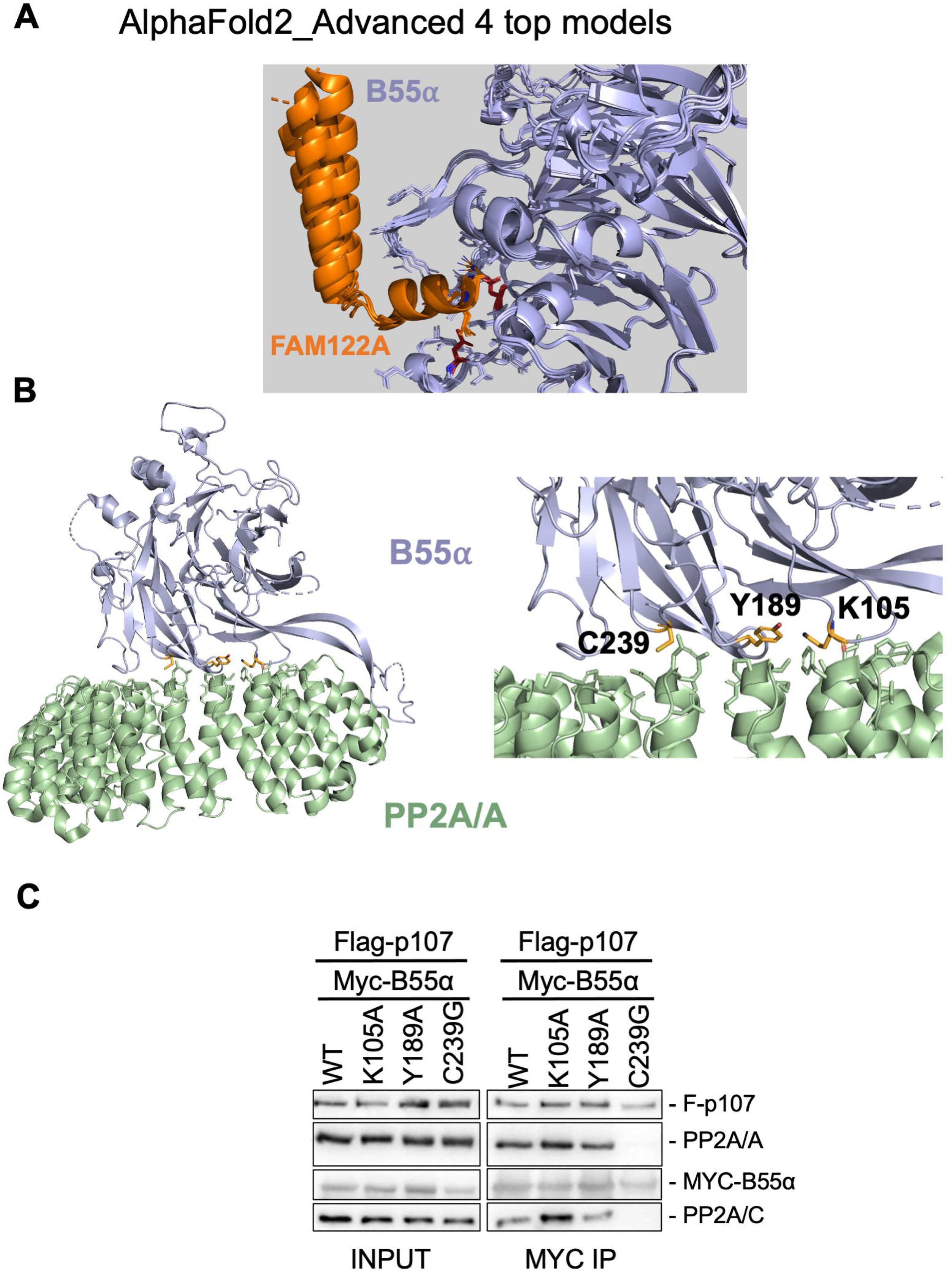
*AlphaFold2_advanced* top models and B55α mutants that render it monomeric used in proteomic analyses. **A.** *AlphaFold2_advanced* prediction models (top 4) of FAM122A binding to B55α **B.** The PP2A/B55α interface with B55α sites predicted to contact PP2A/A. **C.** B55α mutants were assayed for binding to the A-C dimer via co-transfection followed by immunoprecipitations and western blot analysis.

**Suppl. Fig. 5.**
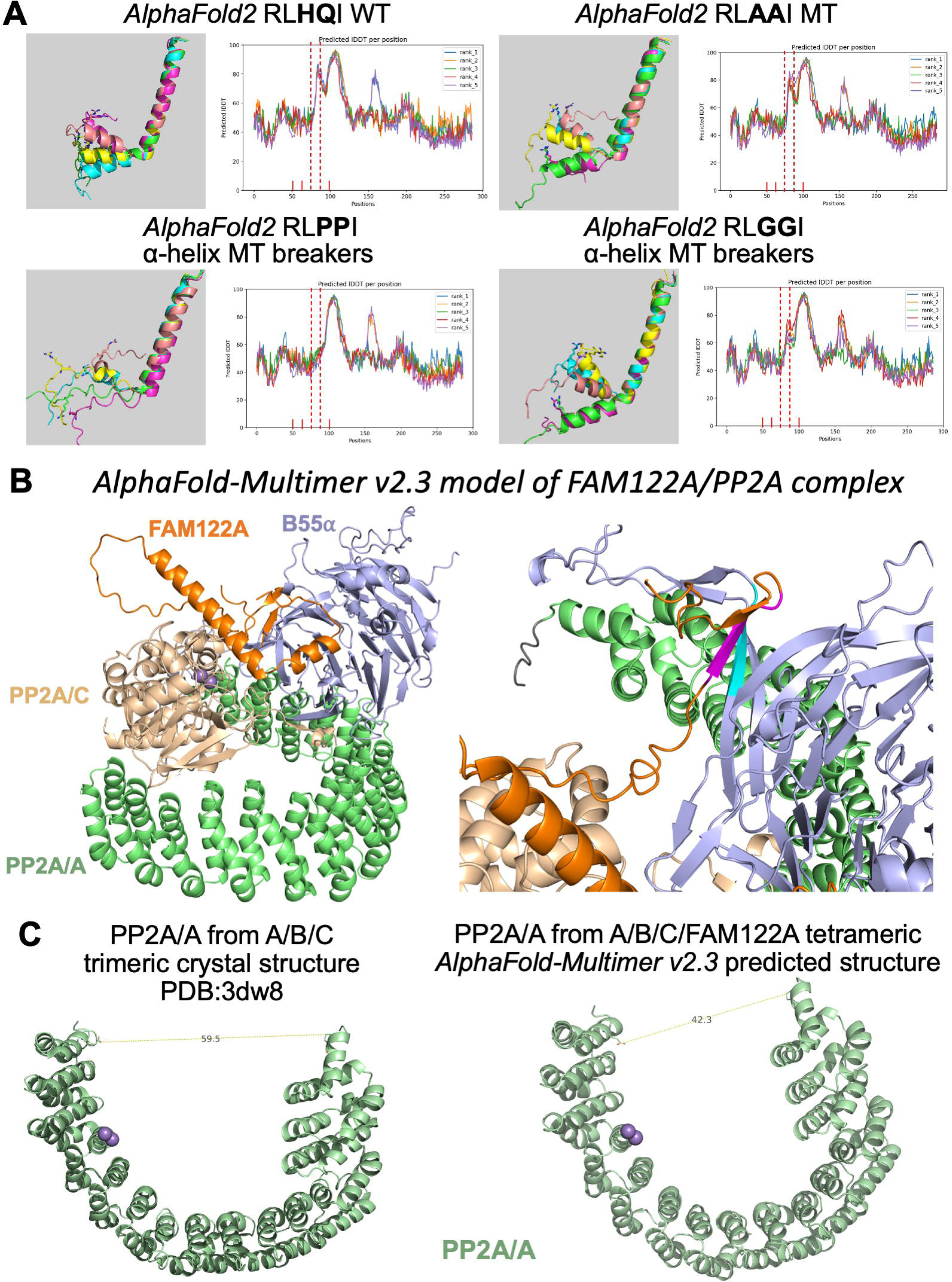
Mutation of amino acids in helix-1 not involved in specific contacts with B55α to Pro or Gly but not Ala are predicted to break or distortion helix-1. **A.** ColabFold (*AlphaFold2_advanced*) prediction models and scores for FAM122A WT and mutations to disrupt helix formation. The region of helix-1 region is flanked by red-dashed lines demonstrating lower structural confidence scores. **B.** Cartoon model of the FAM122A/PP2A/B55α holoenzyme generated in AlphaFold_Multimer v2.3. **C.** Comparison of the curvature of the PP2A/A (scaffold) subunit from the 3dw8 (59.5 Å) crystal structure to the FAM122A/PP2A/B55α tetrameric complex (42.3 Å).

**Suppl. Fig. 6.**
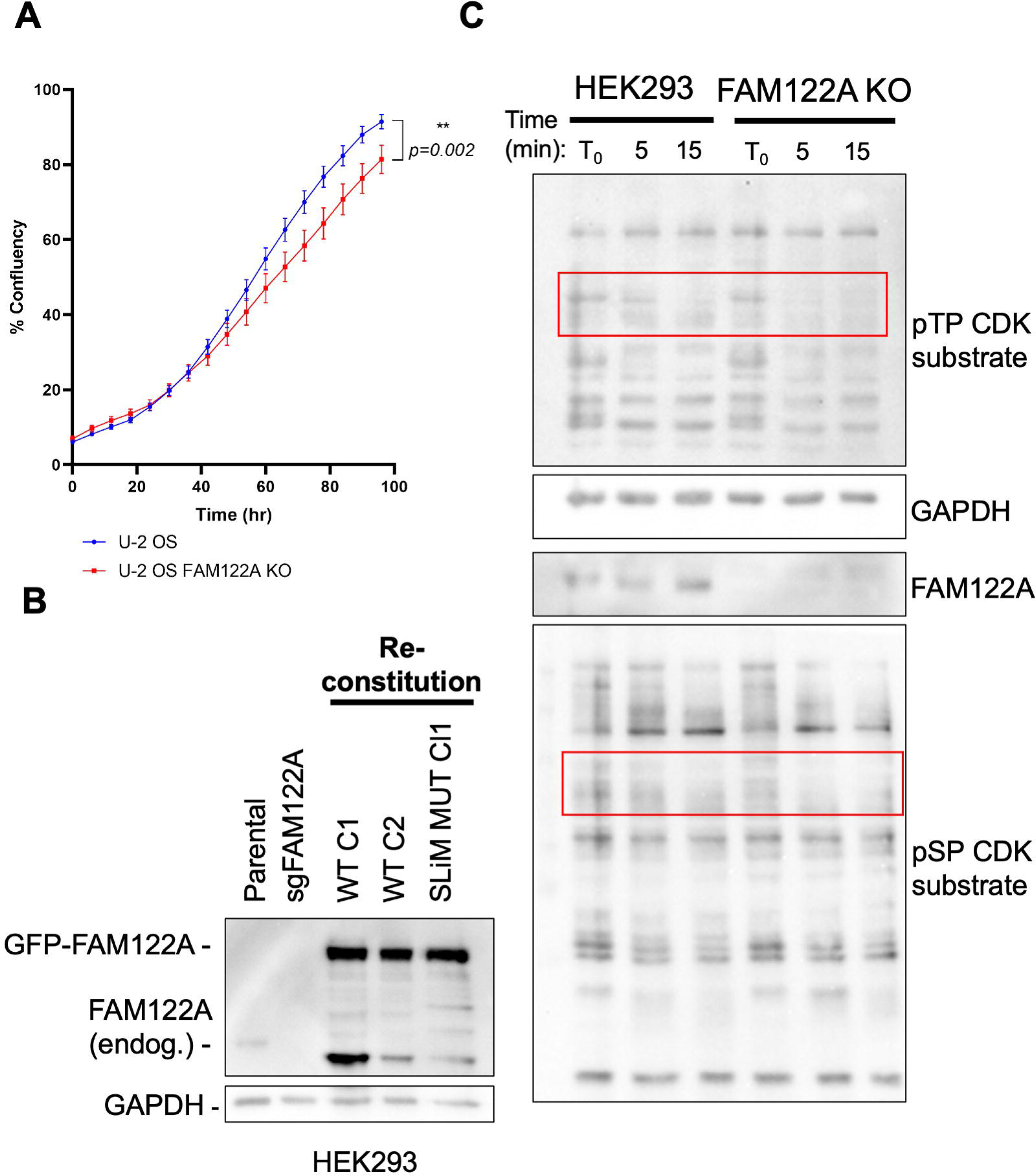
FAM122A Knockout in U2-OS cells, HEK293 FAM122A reconstitution and substrates phosphorylated on CDK SP/TP in protein lysates lacking FAM122A are dephosphorylated with faster kinetics. **A.** Proliferation curves of FAM122A KO in U2-OS cells display similar proliferative defects. **B.** Western blot for FAM122A in HEK293 cells, KO and WT/MT reconstitution. **C.** Western blot for phospho-CDK substrates (TP and SP) to demonstrate phosphatase activity in FAM122A expressing versus KO cells. Cells were lysed in the absence of phosphatase inhibitors with inhibitors being added at indicated timepoints.

**Suppl. Fig. 7.**
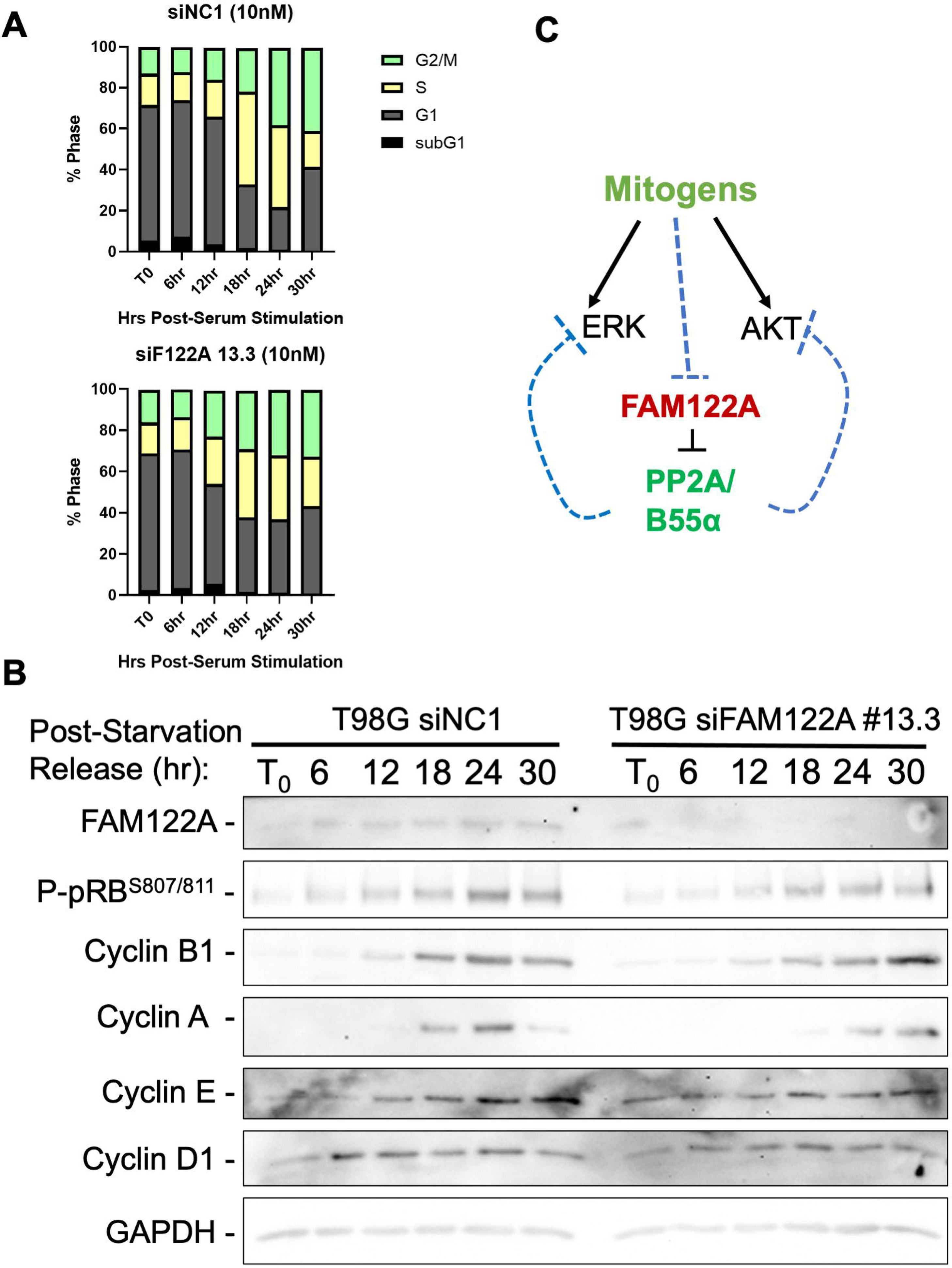
Knockdown of FAM122A with siRNA in serum starved T98G cells delays progression through G1/S, pRB phosphorylation and expression of cyclins. **A.** PI cell cycle analysis of FAM122A knockdown in T98G cells shows delayed cell cycle progression. **B.** Western blot analysis detailing delays in pRB phosphorylation and expression of cyclins. **C.** Pathway schematic demonstrating the role of FAM122A on B55α attenuation of ERK and AKT activity.

**Suppl. Fig. 8.**
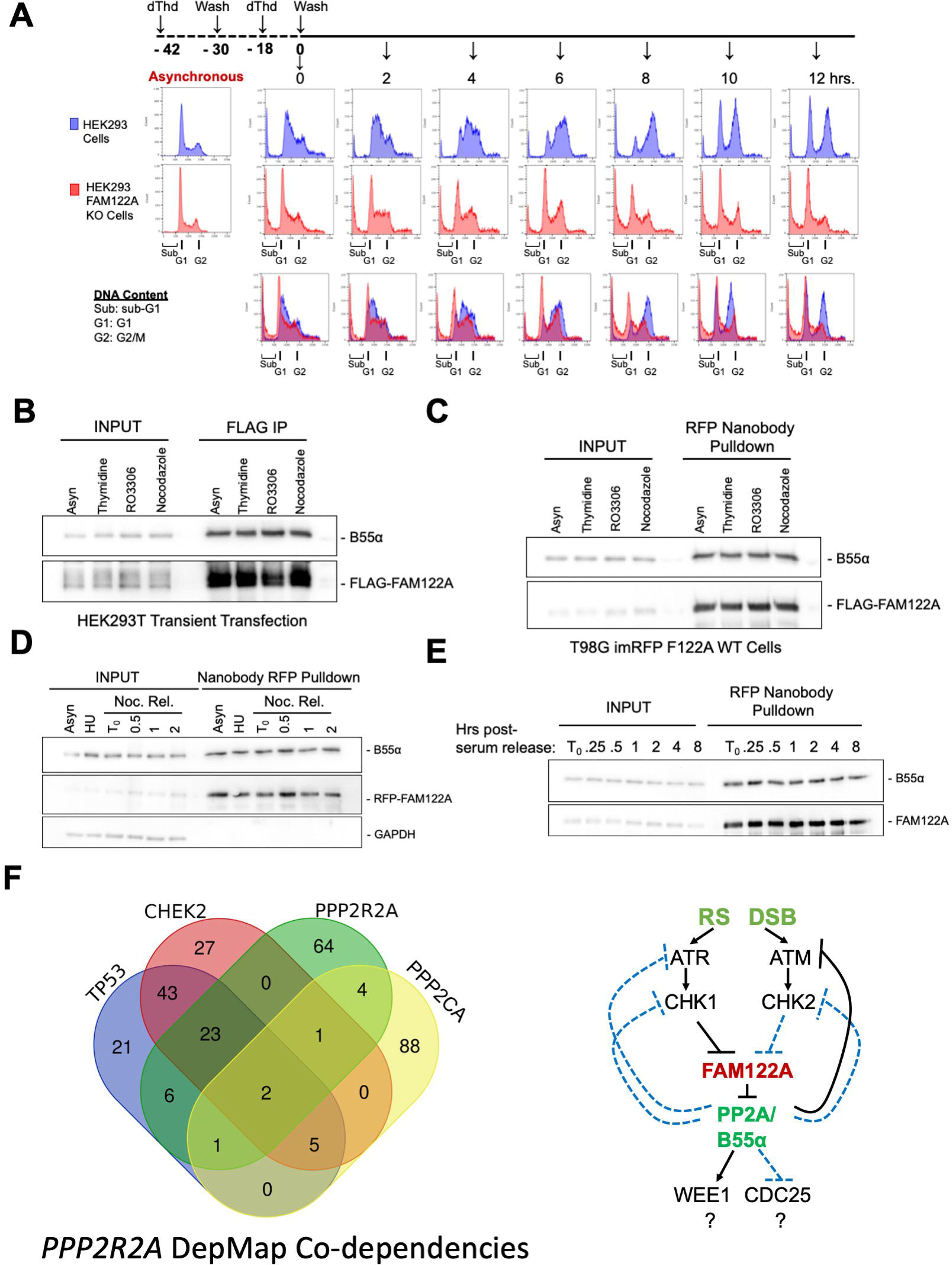
Double thymidine block release of FAM122A KO cells and FAM122A is constitutively bound to B55α during the cell cycle and the G1/S and intra-S checkpoints. **A.** Schematic and cell cycle PI profiles of HEK293 and FAM122A KO cells released from a double thymidine block. FAM122A KO cells fail to synchronize like parental HEK293 cells and exhibit increased SubG1 DNA content. **B-C.** Western blot of pulldowns of FLAG-FAM122A HEK293T and reconstituted RFP-FAM122A T98G cells treated for 24 hr with DMSO, 2 mM Thymidine, 10 µM RO3306, or 10 nM nocodazole and probed for B55α binding. **D.** Western blot of RFP-nanbody pulldown of RFP-FAM122A T98G cells treated for 24hr with DMSO, 2mM HU, or 10 nM nocodazole. Nocodazole treated cells were also released at given timepoints and probed for B55α binding. **E.** Western blot of RFP-nanbody pulldown of reconstituted RFP-FAM122A T98G cells serum-starved for 72 hrs at indicated points of serum restimulation and probed for B55α binding. **F.** Venn diagram showing common hits among DepMap co-dependencies for *PPP2R2A* (B55α), *CHEK2, TP53, PPP2CA* and a pathway schematic of FAM122A and B55α that involves regulators of the replication stress response.

**Suppl. Fig. 9.**
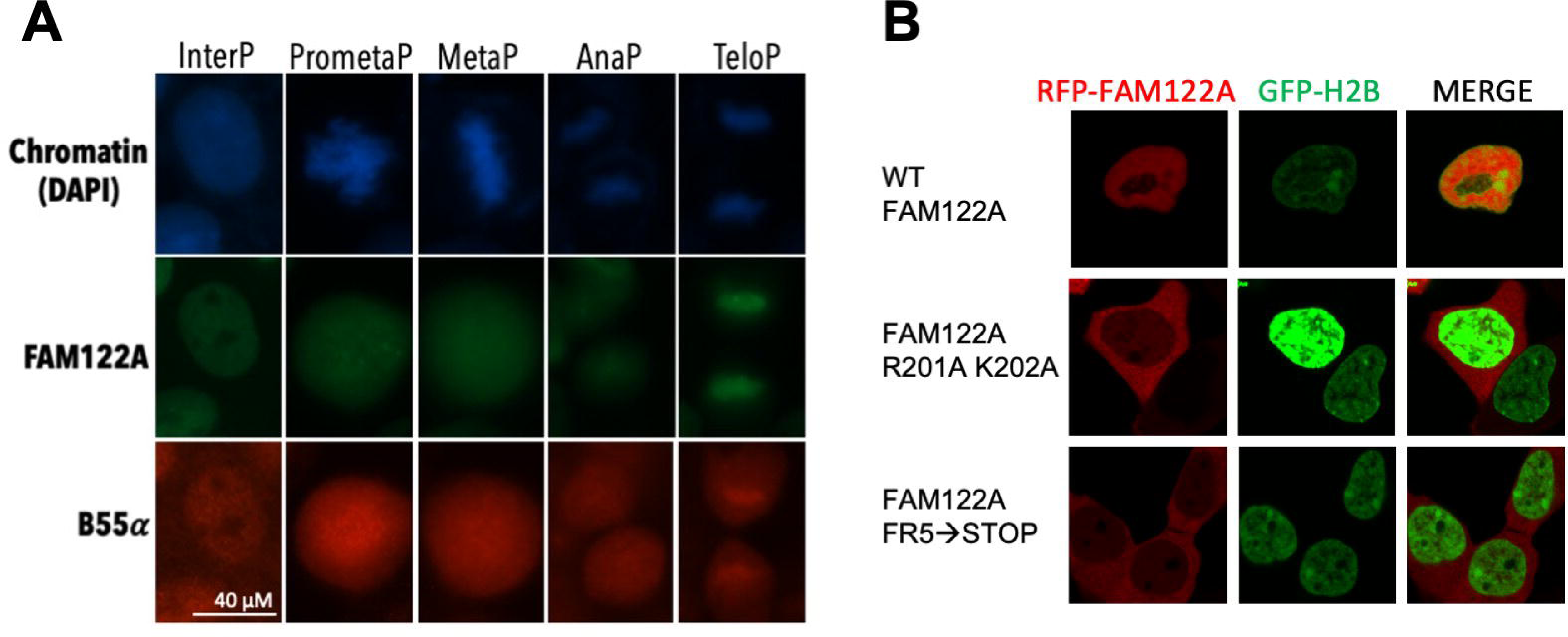
FAM122A is a nuclear protein in interphase that colocalizes with B55α. **A.** Immunofluorescence in U2-OS cells shows that FAM122A and B55α colocalize in the nucleus in interphase, but this is lost in prometaphase (chromosome condensation) and restored late in anaphase. Chromatin was visualized with DAPI and FAM122A and B55α with antibodies. **B.** mRFP-FAM122A WT and NLS deficient MT constructs were co-transfected with GFP-h-H2B to determine the dependency of FAM122A on the proposed NLS in 293 cells. (63x oil immersion confocal images, 5x zoom factor).

